# Resistance bioassays and allele characterisation inform analysis of *Spodoptera frugiperda* (Lepidoptera: Noctuidae) introduction pathways in Asia and Australia

**DOI:** 10.1101/2022.02.08.479273

**Authors:** W. T. Tay, R. V. Rane, W. James, K. H. J. Gordon, S. Downes, J. Kim, L. Kuniata, T. K. Walsh TK

**Affiliations:** CSIRO Black Mountain Laboratories, Clunies Ross Street, ACT 2601, Australia; CSIRO, 343 Royal Parade, Parkville, VIC 3052, Australia; Applied BioSciences, Macquarie University, Sydney NSW 2100, Australia; CSIRO McMaster Laboratories, New England Highway, Armidale NSW 2350, Australia; College of Agriculture and Life Science, Kangwon National University, Republic of Korea; Ramu Agri Industries Ltd., PNG

**Keywords:** FAW, whole genome sequencing, agricultural biosecurity, population genomics, invasion biology

## Abstract

The fall armyworm (FAW) *Spodoptera frugiperda* is present in over 70 countries in Africa, Asia, and Oceania. Its rapid dispersal since 2016 when it was first reported in western Africa, and associated devastation to agricultural productivity, highlight the challenges posed by this pest. Currently, its management largely relies on insecticide sprays and transgenic *Bacillus thuringiensis* toxins, therefore understanding their responses to these agents and characteristics of any resistance genes enables adaptive strategies. In Australia, *S. frugiperda* was reported at the end of January 2020 in northern Queensland and by March 2020, also in northern Western Australia. As an urgent first response we undertook bioassays on two Australian populations, one each from these initial points of establishment. To assist with preliminary sensitivity assessment, two endemic noctuid pest species, *Helicoverpa armigera* and *Spodoptera litura*, were concurrently screened to obtain larval LC50 estimates against various insecticides. We characterised known resistance alleles from the VGSC, ACE-1, RyR, and ABCC2 genes to compare with published allele frequencies and bioassay responses from native and invasive *S. frugiperda* populations. An approximately 10x LC50 difference for indoxacarb was detected between Australian populations, which was approximately 28x higher than that reported from an Indian population. Characterisation of ACE-1 and VGSC alleles provided further evidence of multiple introductions in Asia, and multiple pathways involving genetically distinct individuals into Australia. The preliminary bioassay results and resistance allele patterns from invasive *S. frugiperda* populations suggest multiple introductions have contributed to the pest’s spread and challenge the axiom of its rapid ‘west-to-east’ spread.

## Introduction

The fall armyworm (FAW) *Spodoptera frugiperda* (J. E. Smith), is a noctuid moth pest made up of two morphologically indistinguishable strains (C- and R-strains) that is native to tropical and subtropical regions of the Americas. It is highly polyphagous and feeds on host species from at least 76 plant families, principally Poaceae (106 spp.), Asteraceae (31) and Fabaceae (31) (Montezano et al. 2018). In the Americas, it causes significant economic damage to maize, rice, sorghum, millet, soya bean, wheat, alfalfa, cotton, turf, and fodder crops (Nagoshi et al. 2019a). Recognised globally as one of the top 20 arthropod pests (Willis 2017), *S. frugiperda* was first officially confirmed outside of its native range in Western Africa in early 2016 (Goergen et al. 2016) and then detected across virtually all of Sub-Saharan Africa by February 2018 (Nagoshi et al. 2018, Otim et al. 2018, Botha et al. 2019). In July 2018, it was confirmed in Yemen and India, and by December 2018 in Bangladesh, Sri Lanka, Myanmar and Thailand, followed closely by China in January 2019. Nepal confirmed its presence in May 2019, and in July 2019 it was also reported in Egypt, Indonesia, Laos, Cambodia, Malaysia, Vietnam, Taiwan, The Republic of Korea, and Japan (<u>FAO 2021</u>). The moth’s strong flight ability (Rose et al. 1975, Sparks 1979, Jones et al. 2019), its potential to contaminate certain commodities, and ability to act as a ‘hitchhiker’ in trade (Early et al. 2018) contribute to its rapid spread. Whole genome analyses of invasive *S. frugiperda* populations from various African nations, India, China, Southeast Asia, and Australia also suggested that a high proportion of *S. frugiperda* individuals were hybrids of the C- and R-strains (Gui et al. 2020, Tay et al. 2021, 2022a; Zhang et al. 2020, Schlum et al. 2021, Rane et al. 2022).

Much of South and Southeast Asia, and much of Northern Australia, are highly suitable climatically for *S. frugiperda* year-round (du Plessis et al. 2018, Early et al. 2018). It was confirmed in Queensland (Qld; Bamaga Northern Peninsula Area), Australia following clearance of lure traps on 31 January 2020, followed by reports also in north-Western Australia (WA; Kununurra) and in the Northern Territory (NT) in March 2020. Within two weeks of the Bamaga detection, there were reports of *S. frugiperda* damaging maize fields in Strathmore (Qld; *ca*. 1,000 km from the initial Bamaga detection site), followed by successive southward detections of *S. frugiperda* in Qld, marking the pest’s south-eastward antipodean expansion into New South Wales (NSW) in September 2020. Through CLIMEX simulation analysis based on irrigation patterns and rainfall records, *S. frugiperda* is expected to undertake seasonal migrations and may reach as far south as Victoria and Tasmania (du Plessis et al. 2018). Indeed, *S. frugiperda* was reported in Victoria in December 2020 and in Tasmania in April, 2021. Concurrently, and from the Western Australia Kununurra detection, *S. frugiperda* followed the CLIMEX simulated southward expansion patterns along the WA’s western coast, with larval populations detected in the Gingin area (< 80 Km north of the city of Perth) by February 2021.

The incursion of *S. frugiperda* into Australia is believed to have occurred through natural dispersal across the Torres Strait/Timor Sea into northern Australia via a single-entry point (Jing et al. 2021, Qi et al. 2021). However, incursion pathways to Australia could also involve multiple entry points as reported for the biting midges *Culicoides bravitarsis* (Tay et al. 2016), other anthropogenic routes such as infested commodities in trade, as well as through seasonal or on-going new arrivals of individuals from populations that have successfully established in Asia and Southeast Asia (SEA). The eastward spread of *S. frugiperda* across sub-Saharan Africa, the Middle East, the Indian sub-continent, SEA, China, and the Far East (South Korea, Japan) mostly followed chronological detection and confirmation which led to a widely accepted belief that an invasive population established in west Africa (Goergen et al. 2016) through a single founder, similar to the ‘invasive bridgehead effect’ (Guillemaud et al. 2011), was the starting point for its subsequent global expansion. This hypothesis is supported by an overwhelmingly homogeneous genetic signature based on single partial gene markers (e.g., Cock et al. 2017, Nagoshi et al. 2018).

Whole genome analysis and genome-wide single nucleotide polymorphic marker analysis of invasive populations from Africa (Benin, Uganda, Malawi, Tanzania) and Asia (India, China) instead suggest multiple founding events (Tay et al. 2021, 2022a; Zhang et al. 2020, Schlum et al. 2021); a similar conclusion reached also from analysis of multiple *S. frugiperda* populations from across China based on microsatellite DNA markers (Jiang et al. 2022). Recently, population genomic analysis of *S. frugiperda* populations from SEA (i.e., Philippines, Vietnam, Laos, Malaysia, Myanmar), from East Asia (i.e., South Korea), and Pacific/Oceania (i.e., Papua New Guinea; Australia) identified significant genetic diversity in these invasive *S. frugiperda* populations, and distinct population signatures between populations in close proximity (e.g., Kedah and Penang populations from Malaysia, and between Qld, WA and NT populations in Australia (Rane et al. 2022); between populations within e.g., Yunnan province (Tay et al. 2022a), and populations from Anhui and Jiangsu (Jiang et al. 2022)). Such distinct population genomic structure contradicted the expected gene flow signatures of a single introduction and a west-to-east spread of this pest. Instead, they suggested relatively limited and localised spread of populations while also identified the likely multiple independent introduction pathways of *S. frugiperda* into the region including in SEA, Asia and Australia (Rane et al. 2022).

Distinguishing among single vs. multiple introductions within Australia (and elsewhere), and understanding the resistance profiles of existing and any new incursions, is therefore critical for informing the future management of this pest. For example, this information will assist with forecasting likely resistance profiles of invasive populations selected outside of Australia, and could prioritise efforts to prevent the introgression (potentially of multiple separate introductions) of known insecticide resistance genes and alleles from endemic populations (e.g., Carvalho et al. 2013, Banerjee et al. 2017, Boaventura et al. 2020a, Guan et al. 2020) into invasive populations, or through reciprocal introgression of newly selected/developed resistance traits from invasive populations to native populations.

In this study, we report on the first bioassay experiments and resistance gene characterisation that aimed to understand how the initial invasive populations of *S. frugiperda* in Australia responded to selected approved insecticide compounds and *Bacillus thuringiensis* (Bt) toxins in commercial transgenic plant varieties and available as sprays for the horticultural, grains and cotton industries. Since populations were presumed collected before selection occurred in Australia, these responses served as the first indication of selection against insecticides and Bt toxins used elsewhere (i.e., in the species’ native range and in its recent invasive ranges from Southeast Asia, Asia, and/or Africa (Schlum et al. 2021; Tay et al. 2022a; Jiang et al. 2022; Rane et al. 2022)) prior to arrival. While sampling from multiple Australian regions would be ideal (but impossible at the time as our laboratory had only been provided with these two lines during the pandemic travel restriction), comparing the response differences between these two *S. frugiperda* populations from different northern Australia regions nevertheless could demonstrate the use of bioassay data to distinguish signatures of possible multiple points of entry compared with a single entry and spread event as generally assumed (e.g., Jing et al. 2021, Qi et al. 2021). To provide a first insight into the efficacy of key compounds and toxins we compared the responses of *S. frugiperda* with two related major endemic crop pests in Australia, *Helicoverpa armigera* (subspecies *conferta;* Hardwick 1965, Anderson et al. 2016, Pearce et al. 2017b, a, Zhang et al. 2022) and *S. litura*, which are currently managed through adaptive resistance management plans. Finally, we provided a critical review of resistance profiles in selected invasive populations of *S. frugiperda* globally to better understand its invasion biology, and consider our work alongside a subsequent more comprehensive examination in Australia of insecticide resistance profiles (Bird et al. 2022). Our study is the only available information for Australian populations on the vulnerability of new arrived *S. frugiperda* to relevant Bt toxins.

## Material and Methods

### Live insects

Live *S. frugiperda* populations were sourced from Queensland (Qld) and Western Australia (WA). This population (CSIRO code: *‘S*f20-1’ from Qld; see Table 2) consisted of 30 field-collected pupae (of which 29 pupated and emerged as adults) for setting up the laboratory colony, was first noticed around 19^th^ Feb, 2020, was collected on 3^rd^ March, 2020, and represented the first reported *S. frugiperda* detected attacking maize in Australia’s agricultural landscape. The sampling site for this population was from the University of Queensland (UQ) field station (Rex Road, Walkamin Qld 4872) Strathmore Station, (Lat/Lon: -17.17892, 145.43359, Elevation 685m).

A second *S. frugiperda* population (CSIRO code: ‘Sf20-4’ from WA; see Table 2) was collected as larvae from a maize crop from the Northern Australian Crop Research Alliance Pty Ltd, Kununurra, Western Australia. These larvae were collected on 17-Aug, 2020 from the Kununurra Airport field trail block (off Victoria Highway; - 15.10090, 128.81342) on V4-6 growth stage maize, prior to applications of insecticides. A total of 30 larvae of different instar stages arrived (= F_0_), and 10 (six females and four males) survived to pupate and emerge as adult moths to produce >200 F_1_ individuals and to generate sufficient larvae to commence bioassays from F_4_.

Two related endemic noctuid species, *S. litura* and *Helicoverpa armigera conferta* (hereafter ‘*H. armigera’*), that are pests of a range of broadacre cropping and horticulture industries (and especially grains and cotton) were also included in the bioassay experiments to assist with interpretating results such as base line susceptibility and tolerance levels to the chemicals and toxins tested. Approximately 30 *S. litura* larvae with unknown resistance profiles (from Mareeba and Toowoomba, Qld) pupated and were used to start the laboratory colony. The *H. armigera* colony (CSIRO general rearing ‘GR’ strain) is a laboratory strain housed at the Black Mountain Laboratory in the ACT that was established during the mid-1980’s with individuals originally collected as eggs from cotton fields around Narrabri NSW. Initially the GR colony was periodically replenished to sustain population genetic diversity and minimise inbreeding but no new material has been introduced for around two decades and the colony is susceptible to many chemical insecticides, including organochlorines, organophosphates, carbamates and pyrethroids, as well as to the Cry1Ab, Cry1Ac, and Cry2Ab Bt toxins (Pearce et al. 2017a, Pearce et al. 2017b). It therefore has tolerance levels that are not dissimilar to those of a progenitor population created in 2011 that was deliberately maintained as susceptible through screening (see Bird et al. 2021).The *H. armigera* GR population was confirmed by bioassays as having no Cry1Ac/Cry2Ab/VIP3A resistance alleles. The field collected *S. frugiperda* strains were tested within 4 generations of establishment in the laboratory.

### Colony maintenance

*Helicoverpa armigera* and *Spodoptera* species (*S. frugiperda* Sf20-1 and Sf20-4; *S. litura)* were reared by the method of Teakle and Jensen (1985) at 25 °C, 50 ± 10% relative humidity, and with a light/dark cycle of 14:10 h. Artificial diet for *H. armigera* consisted of 81 g soya flour, 37.5 g wheat germ, 33 g brewer’s yeast, 2 g ascorbic acid, 2 g nipagen, 3 mL vegetable oil, and 13.5 g agar, which was processed in a microwave oven and made up with water to 1 litre. Diet was poured into rearing cups or bioassay plates, allowed to cool and then stored in the refrigerator (4-8°C) for no longer than 3 days. Diet for *S. frugiperda* was made to 1200mL with water, and contained 100g navy bean flour, 25g soy flour, 60g wheat germ, 30g Brewer’s yeast, 15g casein, 3g Nipagen, 1.5g sorbic acid, 10g vitamin mixture, 4mL vegetable oil, and 18g agar. The mixture was processed in a microwave oven prior to being poured into rearing cups/bioassay plates and allowed to cool as described above for the *H. armigera* diet.

### Bioassays

The bioassay experiments involved commercially available insecticidal compounds and Bt toxins (available in transgenic varieties of cotton and as commercial sprays), as well as Bt toxins that were produced and purified by CSIRO (Table 1). For all insecticides and Bt toxins, Sf20-1 and Sf20-4 *S. frugiperda* populations were tested. The *H. armigera ‘*GR’ line was tested alongside *S. frugiperda* against all chemicals and Bt toxins except Cry1F because it is known to be insensitive to it. We included *S. litura* in all Bt toxin bioassays as it is regarded as a pest of interest by the Australian cotton industry who depend on Bt varieties.

**Table 1:**
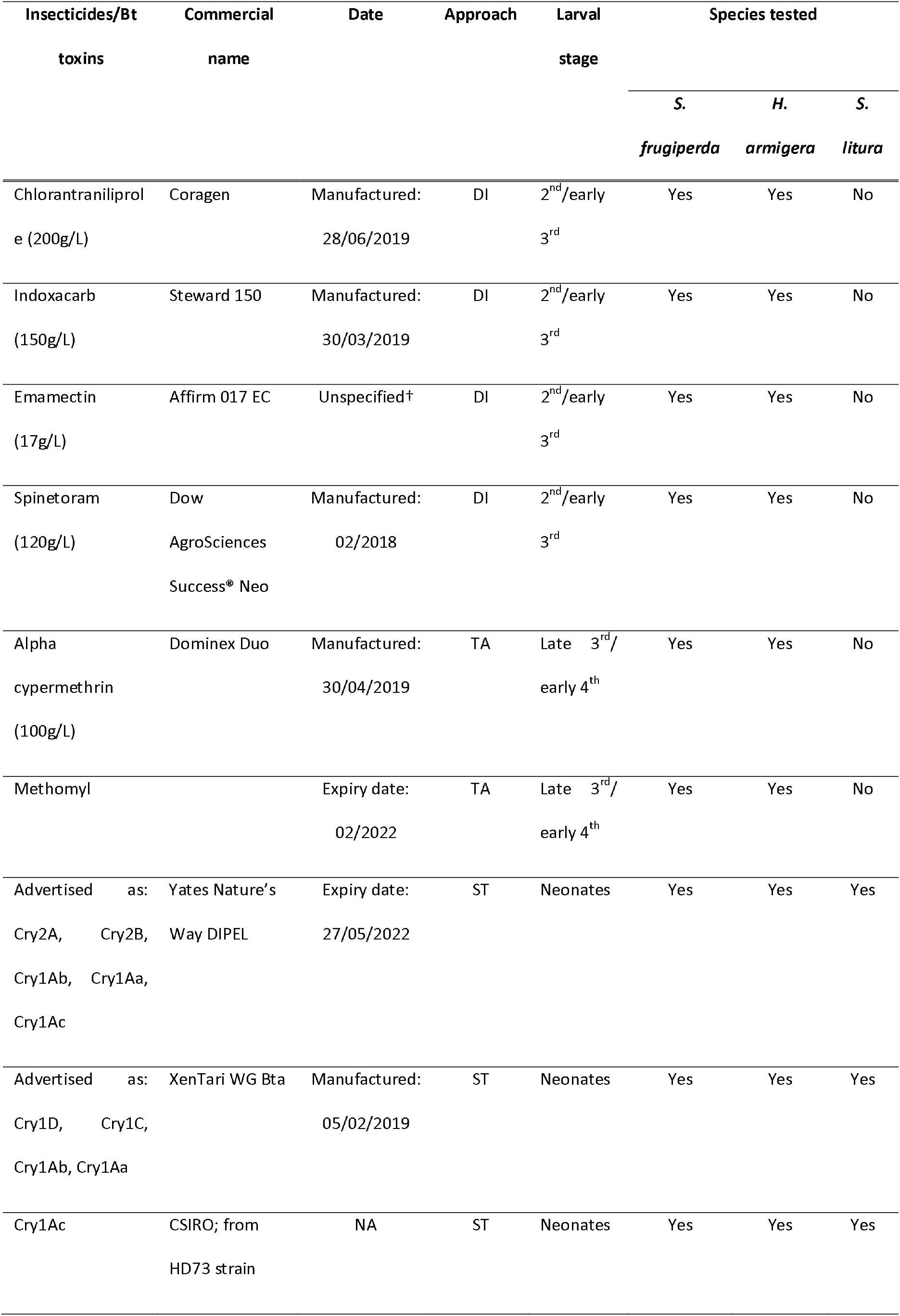
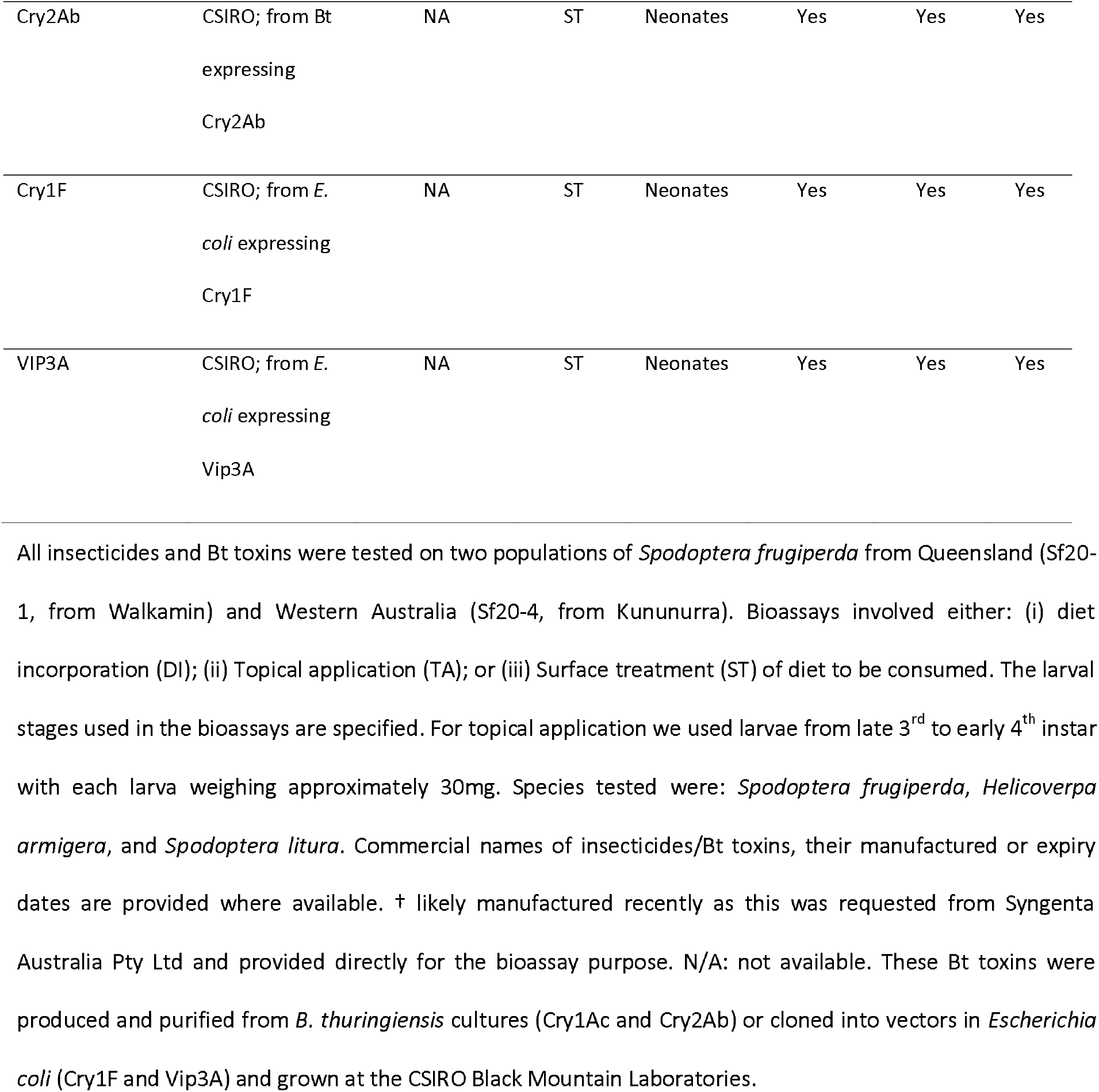
List of insecticides and Bt toxins used in the bioassay experiments.

All bioassays included a negative control for background mortality (i.e., to enable natural mortality rates to be adjusted accordingly) which was treated with the diluent of the individual pesticide. An initial experiment using 10x serial dilutions was performed to narrow the range for the detailed bioassay using 8-16 individuals per dose replicated once. The full bioassay involved 2x serial dilutions at 6-7 doses with 16-24 individuals per dose replicated 3-5 times. The bioassays involved diet incorporation, surface overlay, or topical application of the insecticidal compounds or Bt toxins (Table 1). The selection among these exposure methods for an insecticide was determined by its mode of entry (contact or ingestion) and the experience of similar bioassays in related noctuid pests (i.e., *H. armigera* and *H. punctigera;* Bird 2015; Bird & Walker 2019).

The surface overlay of Bt toxin assays followed the approach outlined in Mahon et al. (2007b, 2008, 2010, 2012) and Walsh et al. (2014). They were conducted in 96-well trays where each well contained approximately 300µl of rearing diet with a surface area of around 0.567 cm^2^ per well. When the diet cooled, 20µl solution containing an appropriate concentration of toxin was added and allowed to air dry. One neonate was added to each well and the tray was heat sealed with a perforated lidding material.

For diet incorporation, commercial grade insecticides were diluted to the appropriate concentration with water, added to 150ml of artificial diet and mixed well to produce a homogenous solution. Insecticide-incorporated diet was then dispensed into 45-well bioassay trays, each with approximately 1.5⍰ml of diet-insecticide mix. One late-second or early-third instar larva was added to each well and the tray was heat sealed with a perforated lidding material (as per Bird 2015).

Topical bioassays of known concentrations were conducted for alpha-cypermethrin (group 3A) and methomyl (group 1) and involved applying drops of pesticide as described in Bird (2018). Specifically, 1 μl of acetone/insecticide solution was applied to the dorsal thorax of 3^rd^ to 4^th^ instar larvae (30-40g) using a 50 μl micro-syringe. Bioassay trays were incubated at 25 °C, 45-55% RH, and a photoperiod 14:10 (L:D) h for 6 days (e.g., see Bird and Akhurst 2007, Downes et al. 2009). The numbers of dead (moribund; incapable of movement when prodded) and alive larvae (capable of coordinated movement when prodded) were counted and the instars of surviving larvae recorded. The LC_50_ for each toxin was calculated from pooled raw data by probit analysis using the POLO-PC program (LeOra Sorftware 1987).

### Spodoptera frugiperda specimens for genomic analyses

Populations of *S. frugiperda* from South Korea (SK), Papua New Guinea (PNG), and Peru were included for whole genome sequencing (Table 2). The methods for SK and PNG specimen preservation involved collection from fields and placing larvae directly into high concentrations of ethanol (95-99.9%) to transport to the laboratory where they were then stored at -20°C and replaced with fresh ethanol after 24-48 hours. Peru specimens were samples intercepted from Australian pre-border inspections of imported agricultural/horticultural commodities between 2016 and 2019 (see Tay et al. 2021, 2022a). Australian samples (Sf20-1 from Qld, Sf20-4 from WA) were F_0_ individuals and represented individuals obtained directly from fields. Samples from SK, PNG, and Peru were sent to CSIRO and stored at -20°C until DNA extraction.

**Table 2:** *Spodoptera frugiperda* samples from Australia, Papua New Guinea, South Korea, and Peru used in whole genome sequencing for resistance gene characterisation.

| Country | State/Province | Population | N | Date | Note | R : C |
| --- | --- | --- | --- | --- | --- | --- |
| Australia | WA | Kununurra | 9 | 16-Aug-20 | 'Sf20-4', represent the original WA field-collected individuals | 9 : 0 |
|  | Qld | Strathmore Station, Walkamin | 29 | 03-Mar-20 | 'Sf20-1', represent the original Qld field-collected individuals | 29 : 0 |
| South Korea | Milyang | MF | 7 | Feb-20 | Milyang-si, Gyeongsangnam-do, G<sub>4</sub> lab, GPS: 35°29.29 N, 128°44.31 E | 7 : 0 |
|  | Goryong | GR | 1 | Sep-19 | Goryeong-gun, Gyeongsangbuk-do, Corn-field; GPS: 35° 64.04 N, 128°39.08 E | 1 : 0 |
|  | Haenam | MN | 1 | Aug-19 | Haenam-gun, Jeollanam-do, Corn field; GPS: 34°24.38 N, 126°37.55 E | 1 : 0 |
|  | Milyang | MY | 2 | Sep-19 | Milyang-si, Gyeongsangnam-do, Corn field; GPS: 35°29.29 N, 128°44.31 E | 0 : 2 |
|  | Muan | MA | 1 | Aug-19 | Muan-gun, Jeollanam-do, Corn field; GPS: 34°52.21 N, 126°31.11 E | 1 : 0 |
| Papua New Guinea | Madang Province | Ramu Sugar Estate | 16 | 15-17 Jun-2020 | 5°58.154 S, 145°53.252 E, Maize host | 16 : 0 |
|  | Central Province | Yule Island Junction | 1 | 5-Jun-20 | corn host | 1 : 0 |
|  |  | Pre-border intercepted specimens on imported agricultural |  |  |  |  |
| Peru | n/a | commodities | 16 | 2016 - 2019 | Samples from Tay et al. (2020) (Suppl. Table 1, individuals PE01 – PE16) | 0 : 16 |

Australian *S. frugiperda* populations were Sf20-1 from Qld and Sf20-4 from WA. Number of specimens from each location is indicated by 'N'. Where available, GPS co- ordinates and elevation are provided. The number of individuals belonging to the *S. frugiperda* R-strain or the C-strain mitochondrial DNA COI haplotypes is also provided.

### DNA extraction and genome library preparation

DNA of individual *S. frugiperda* samples was extracted using the Qiagen DNA extraction kit and eluted in 200µL elution buffer following protocols as provided by the manufacturer (Qiagen, Hilden Germany). Genomic DNA libraries for individual samples were prepared, quantified and sent for commercial sequencing by the Australian Genome Research Facility (AGRF) in Melbourne, Victoria, Australia.

### Processing of genome sequences

Genome sequencing data for individuals were trimmed to remove adapter sequences using trim_galore (v 0.6.6; https://www.bioinformatics.babraham.ac.uk/projects/trim_galore/) and aligned to the *S. frugiperda* rice genome (v1.0) (Gouin et al. 2017) using bwa_mem2 (v2) (Vasimuddin et al. 2019). Duplicate alignments were removed using SAMBLASTER (v 0.1.26) (Faust and Hall 2014) and sorting completed using SAMtools (v1.9) (Li et al. 2009).

### Strain identification and resistance alleles characterisation by whole genome sequencing

For identification of *S. frugiperda R*- or C-strain, the used the partial mtCOI gene sequences for the R-strain (GenBank MF197867) and the C-strain (GenBank MF197868) of Otim et al. (2018) as reference sequence for mapping against the whole genome sequence data for each individual *S. frugiperda* from Sf20-1, Sf20-4, SK, PNG, and Peru. Mapping of partial mtCOI gene was carried out within Geneious v11.1.5 (Biomatters Ltd, Auckland, NZ) using the Geneious Mapper program with assembly parameters specified to ‘Low Sensitivity / Fastest’, with no trimming before mapping, and the ‘Fine Tuning’ option set to ‘Iterate 2 times’ to map reads to the consensus from the previous iteration. Full mitochondrial DNA genomes of all individuals were also assembled following the procedures as outlined in (Otim et al. 2018) and have been reported by Rane et al. (2022). For all resistance alleles of interest (ABCC2, ACE-1, RyR, VGSC; see Table S1), base-sequence at the genomic location for individuals listed in Table 1 was extracted from the alignment file using BCFTOOLS MPILEUP followed by CALL, since BBMAP does not report ‘non-variant’ sites. Results were tabulated for inference.

### Comparisons of published ACE-1 and VGSC resistance allele frequencies and published indoxacarb and chlorantraniliprole LC_50_ values

Three resistance loci for the organophosphate/carbamate ACE-1 gene have been reported to-date in *S frugiperda:* (i) A201S, (ii) G227A, and (iii) F290V. To understand the frequencies of these alleles, we surveyed specimens from invasive (i.e., Australia, PNG, South Korea) and native (Peru) *S. frugiperda* populations, and combined this information with reported allele frequencies for these three loci from other native (Brazil, French Guiana, Mexico Guadeloupe, Puerto Rico, USA) and invasive populations (Benin, Uganda, Kenya, Tanzania, Zambia, Malawi; India, Indonesia, China, Australia) (see Table S2 and references therein). For the pyrethroid *para* sodium channel resistance gene VGSC, three loci have been identified to date in *S. frugiperda:* (Carvalho et al. 2013, Guan et al. 2020, Yainna et al. 2021). Resistance and susceptible allele frequencies in these three VGSC loci were also surveyed from published studies from native and invasive *S. frugiperda* populations (see Table S2).

For comparisons of bioassay LC_50_ data for indoxacarb and chlorantraniliprole between published data and our study, we considered only those that reported broadly similar methodologies (see Table 1) such as clearly stated route of delivery (i.e., diet incorporation/ingestion) for the insecticidal compounds, and using comparable developmental stages of larvae (i.e., 2^nd^/early 3^rd^ instar) and scoring criteria.

## Results

### Bioassays of Individual Bt proteins

Cry1Ac was not highly effective against the *Spodoptera* species tested in this study (Fig. S1; Table 3). Based on the LC_50_, and relative to *H. armigera, S. frugiperda* Sf20-1 was 174x less sensitive, *S. frugiperda* Sf20-4 was 99x less sensitive, and *S. litura* was 120x less sensitive. This suggests that in *S. frugiperda*, Cry1Ac would give similar control in Australia to that found in *S. litura* but far less than for *H. armigera*.

**Table 3:** Summary bioassay data on *Spodoptera frugiperda* populations from Queensland and Western Australia, *S. litura*, and *Helicoverpa armigera* involving surface treatment of the diet with Bt toxins and products.

| Pesticide | Populations/<br>species | N | Slope | LC<sub>50</sub> | 95% CI | LC<sub>99</sub> | 95% CI | $\chi^2$ (Degrees of<br>Freedom) | P | Toxicity ratio (*H.*<br>*armigera* = 1) |
| --- | --- | --- | --- | --- | --- | --- | --- | --- | --- | --- |
| Cry1Ac | *H. armigera* | 741 | 1.837 ±0.126 | 0.025 | 0.021 - 0.029 | 0.465 | 0.316 - 0.773 | 31.66 (29) | 0.334 | - |
| (µg/cm<sup>2</sup>) | Sf20-1 | 504 | 0.827 ±0.102 | 4.34 | 1.81 - 8.15 | 80.6 | 36.85 - 249.17 | 18.38 (19) | 0.497 | 174 |
|  | Sf20-4 | 575 | 1.449 ±0.149 | 2.48 | 1.79 - 3.29 | 85.34 | 53.32 - 159.68 | 17.77 (22) | 0.720 | 99 |
|  | *S. litura* | 501 | 1.626 ±0.135 | 2.99 | 2.06 - 4.20 | 80.7 | 38.60 – 284.54 | 49.14 (19) | <0.001 | 120 |
| Cry2Ab | *H. armigera* | 476 | 1.501 ±0.143 | 0.049 | 0.034 -0.068 | 1.74 | 0.86 - 6.57 | 31.834 (18) | 0.023 | - |
| (µg/cm<sup>2</sup>) | Sf20-1 | 575 | 1.649 ±0.136 | 0.655 | 0.435 - 0.951 | 16.88 | 7.24.- 86.01 | 86.222 (22) | <0.001 | 13 |
|  | Sf20-4 | 551 | 2.201 ±0.246 | 0.178 | 0.138 - 0.221 | 2.03 | 1.33 - 4.94 | 39.076 21 | 0.010 | 4 |
|  | *S. litura* | 859 | 1.197 ±0.081 | 0.511 | 0.335 - 0.734 | 44.9 | 20.179 - 154.39 | 91.946 33 | <0.001 | 10 |
| Cry1F | Sf20-1 | 576 | 1.531 ±0.115 | 0.025 | 0.015 - 0.038 | 0.838 | 0.365 - 3.949 | 87.309 (22) | <0.001 | - |
| (µl/cm<sup>2</sup>) | Sf20-4 | 575 | 1.809 ±0.149 | 0.021 | 0.014 - 0.029 | 0.394 | 0.211 - 1.138 | 19.947 (22) | 0.586 | - |
|  | *S. litura* | 574 | 1.524 ±0.121 | 0.0088 | 0.0069 - 0.011 | 0.294 | 0.187 - 0.535 | 58.968 (22) | <0.001 | - |
| VIP3a | *H. armigera* | 763 | 1.486 ±0.093 | 0.0062 | 0.0051 - 0.0075 | 0.230 | 0.140 - 0.400 | 33.738 (30) | 0.291 | - |
| (µl/cm<sup>2</sup>) | Sf20-1 | 619 | 1.881 ±0.225 | 0.0021 | 0.0010 - 0.0031 | 0.049 | 0.021 - 0.257 | 68.846 (24) | <0.001 | 0.152 |
|  | Sf20-4 | 599 | 2.398 ±0.212 | 0.0019 | 0.0016 - 0.0023 | 0.018 | 0.013 - 0.028 | 20.934 (23) | 0.585 | 0.078 |
|  | *S. litura* | 575 | 1.693 ±0.129 | 0.00065 | 0.0005 - 0.0008 | 0.015 | 0.088 - 0.035 | 39.108 (22) | 0.013 | 0.065 |
| Dipel | *H. armigera* | 526 | 1.832 ±0.154 | 2.11 | 1.59 - 2.72 | 39.2 | 22.43 - 92.76 | 33.878 (20) | 0.007 | - |
| (IU/cm<sup>2</sup>) | Sf20-1 | 644 | 1.832 ±0.154 | 52.02 | 36.55 - 70.05 | 2612.6 | 1379.89 - 6752.99 | 29.258 (25) | 0.253 | 24 |
|  | Sf20-4 | 574 | 1.703 ±0.153 | 38.93 | 23.97 - 56.35 | 904.1 | 469.80 - 2779.63 | 53.767 (22) | 0.002 | 18 |
|  | *S. litura* | 788 | 1.426 ±0.098 | 35.96 | 28.23 - 44.99 | 1540.8 | 916.66 - 3110.88 | 40.916 (31) | 0.110 | 17 |
| XenTari | *H. armigera* | 620 | 1.530 ±0.136 | 5.95 | 4.69 - 7.40 | 197.2 | 118.7 - 398.3 | 23.051 (24) | 0.516 | - |
| (DBM/cm<sup>2</sup>) | Sf20-1 | 647 | 2.375 ±0.289 | 11.81 | 8.33 - 15.17 | 112.7 | 69.8 - 270.1 | 39.18 (25) | 0.035 | 2 |
|  | Sf20-4 | 549 | 1.333 ±0.180 | 19.01 | 9.27 - 29.8 | 1058.9 | 379.5 - 10739.7 | 43.25 (21) | 0.003 | 3 |
|  | *S. litura* | 741 | 2.375 ±0.289 | 24.86 | 17.08 - 31.14 | 206.9 | 126.2 - 597.5 | 47.92 (29) | <0.001 | 4 |

*S. frugiperda* populations were from Walkamin, Queensland (Sf20-1) and Kununurra, Western Australia (Sf20-4); *S. litura* was from Mareeba Queensland, and *Helicoverpa* *armigera* was a laboratory 'GR' line. The concentration of each pesticide required to kill 50% and 99% of the test subjects (*H. armigera*, *S. frugiperda*, *S. litura*) are given as LC<sub>50</sub> (50% lethal concentration) and LC<sub>99</sub> (99% lethal concentration), respectively. 95% confidence intervals (95% CI) for both LC<sub>50</sub> and LC<sub>99</sub> are also provided. Sample sizes (N) of *H. armigera*, *S. litura*, and the two *S. frugiperda* laboratory culture lines used in the bioassay tests are indicated. P-values (*P*) associated with the $\chi^2$ tests are also provided.

The *Spodoptera* species are less sensitive to Cry2Ab than *H. armigera* but the differences are not as striking as for Cry1Ac (Fig. S2; Table 3); *S. frugiperda* Sf20-1 was 13x less sensitive, *S. frugiperda* Sf20-4 was 4x less sensitive and *S. litura* was 10x less sensitive. This suggests that in *S. frugiperda*, Cry2Ab would give similar control in Australia to that found in *S. litura* which is unlikely to be different from *H. armigera* in terms of field control.

Cry1F is a Bt protein that has been deployed in certain genetically modified plants to target *Spodoptera.* Though not effective against *H. armigera sensu lato* even at a relatively high dose, Cry1F was effective against *S. frugiperda* at a much lower dose (Fig S3; Table 3), indicating that this species is much more sensitive to this Bt protein than *H. armigera*. Relative to *S. litura,* the LC_50_ data shows that *S. frugiperda* Sf20-1 and *S. frugiperda* Sf20-4 are 2x less sensitive to Cry1F. In terms of field control there is unlikely to be any distinguishable difference in Cry1F efficacy against *S. frugiperda* vs. *S. litura*.

*S. frugiperda* Sf20-1 and Sf20-4 showed a similar tolerance to Vip3A as *S. litura* (∼2.34x and 1.2x higher, respectively) and all three populations were more sensitive to Vip3A than *H. armigera* (0.15x, 0.08x, and 0.07, respectively). In terms of field control there is unlikely to be any distinguishable difference in efficacy of VIP3A against *S. frugiperda* relative to *H. armigera* and *S. litura* (Fig. S4; Table 3).

Based on published studies which show resistance ratios of >2,500 fold in *H. armigera* and *S. frugiperda* that are homozygous for recessive resistance alleles against Bt toxins (e.g., Mahon et al. (2007a) - Cry2Ab, *H. armigera;* Horikoshi et al. (2016), Cry1F, Cry1A, Vip3A, *S. frugiperda)* it is unlikely that either of the founding *S. frugiperda* populations in Australia carried resistance alleles to Cry1Ac, Cry2Ab or Vip3A in homozygous states.

#### Foliar Bt

The performance against *S. frugiperda* of the sprayable products containing multiple Bt toxins (XenTari and DIPEL) showed both products to differ in efficacies against *H. armigera, S. frugiperda*, and *S. litura*. XenTari with the mixture of Cry1 toxins including Cry1C was formulated to provide *Spodoptera* control (<u>valentbiosciences</u>, last accessed 13-Jan, 2022) but also for cabbage looper and the diamondback moth (DBM), while DIPEL was formulated for broad spectrum caterpillar control (<u>valentbiosciences</u>, last accessed 13-Jan, 2022). Both DIPEL and XenTari are reported to contain different toxin complements but the relative amounts are unclear. DIPEL contains Cry1Ac and Cry2-type toxins and was less effective against both tested *S. frugiperda* populations than XenTari which contains Cry1Aa, Cry1Ab, Cry1C and Cry1D. Relative to *H. armigera, S. frugiperda* Sf20-1 was 24x less sensitive, *S. frugiperda* Sf20-4 was 18x less sensitive and *S. litura* was 17x less sensitive to Dipel. XenTari toxicity ratios relative to *H. armigera* ranged 2 to 4x for *S. frugiperda* Sf20-1, *S. frugiperda* Sf20-4 and *S. litura* (Figs. S5 and S6; Table 3).

#### Conventional pesticides

Cypermethrin is a pyrethroid pesticide and the bioassay results suggest that *S. frugiperda* (i.e., the Sf20-1 and Sf20-4 laboratory cultures) is less sensitive to it than the laboratory strain of *H. armigera* (56x and 145x respectively: Fig. S7; Table 4).

**Table 4:** Summary bioassay data on *Spodoptera frugiperda* populations and *Helicoverpa armigera* (GR laboratory line) involving topical application to the insect with contact insecticides.

| Toxin | Population/<br>Species | N | Slope | LC<sub>50</sub> | 95% C.I. | LC<sub>99</sub> | 95%CI | $\chi^2$ (Degrees of<br>Freedom) | P | Toxicity ratio (*H.*<br>*armigera* = 1) |
| --- | --- | --- | --- | --- | --- | --- | --- | --- | --- | --- |
| Alpha cypermethrin<br>µg/larvae | *H. armigera* | 1328 | 2.849 ±0.140 | 0.0036 | 0.0032 - 0.0041 | 0.023 | 0.018 - 0.032 | 109.84 (57) | <0.001 | - |
|  | Sf20-1 | 675 | 2.399 ±0.154 | 0.201 | 0.171 - 0.239 | 1.88 | 1.31 - 3.06 | 43.903 (28) | 0.028 | 56 |
|  | Sf20-4 | 766 | 2.186 ±0.132 | 0.523 | 0.427 - 0.641 | 6.06 | 3.99 - 10.85 | 67.208 (32) | <0.001 | 145 |
| Methomyl<br>µg/larvae | *H. armigera* | 858 | 0.809 ±0.059 | 0.057 | 0.031 - 0.097 | 43.12 | 10.27 - 53.93 | 135.89 (36) | <0.001 | - |
|  | Sf20-1 | 765 | 1.064 ±0.076 | 0.254 | 0.177 - 0.356 | 39.2 | 15.81 - 156.30 | 72.309 (32) | <0.001 | 4 |
|  | Sf20-4 | 631 | 0.874 ±0.07 | 2.96 | 1.87 - 4.66 | 1363.7 | 380.11 - 10950.47 | 57.160 (26) | <0.001 | 52 |

*S. frugiperda* from Walkamin Queensland and from Kununurra Western Australia are coded Sf20-1 and Sf20-4, respectively. *Helicoverpa armigera* laboratory line is coded as 'GR'. The concentration of each pesticide required to kill 50% and 99% of the test subjects (*H. armigera*, *S. frugiperda*) are given as LC<sub>50</sub> (50% lethal concentration) and LC<sub>99</sub> (99% lethal concentration), respectively. 95% confidence intervals (95% CI) for both LC<sub>50</sub> and LC<sub>99</sub> are also provided. Sample sizes (N) of *H. armigera* and *S. frugiperda* laboratory culture lines used in the bioassay tests are indicated. P-values (P) associated with the $\chi^2$ tests are also provided.

Bioassay results for the carbamate pesticide methomyl are not as repeatable as other toxins with considerable variability between replicates (Fig. S8; Table 4). However, overall *S. frugiperda* Sf20-4 shows 52x less sensitivity to methomyl than *H. armigera* while *S. frugiperda* Sf20-1 shows a 4x lower sensitivity. This difference is consistent with some level of heterogeneity between the first invasive *S. frugiperda* populations in Australia.

The dose response and LC_50_ for indoxacarb (Fig. S9; Table 5) show that *S. frugiperda* Sf20-1 and *S. frugiperda* Sf20-4 are less sensitive to this chemistry than *H. armigera* (22x and 208x respectively). This difference is consistent with some level of heterogeneity between the first invasive *S. frugiperda* populations in Australia.

**Table 5:**
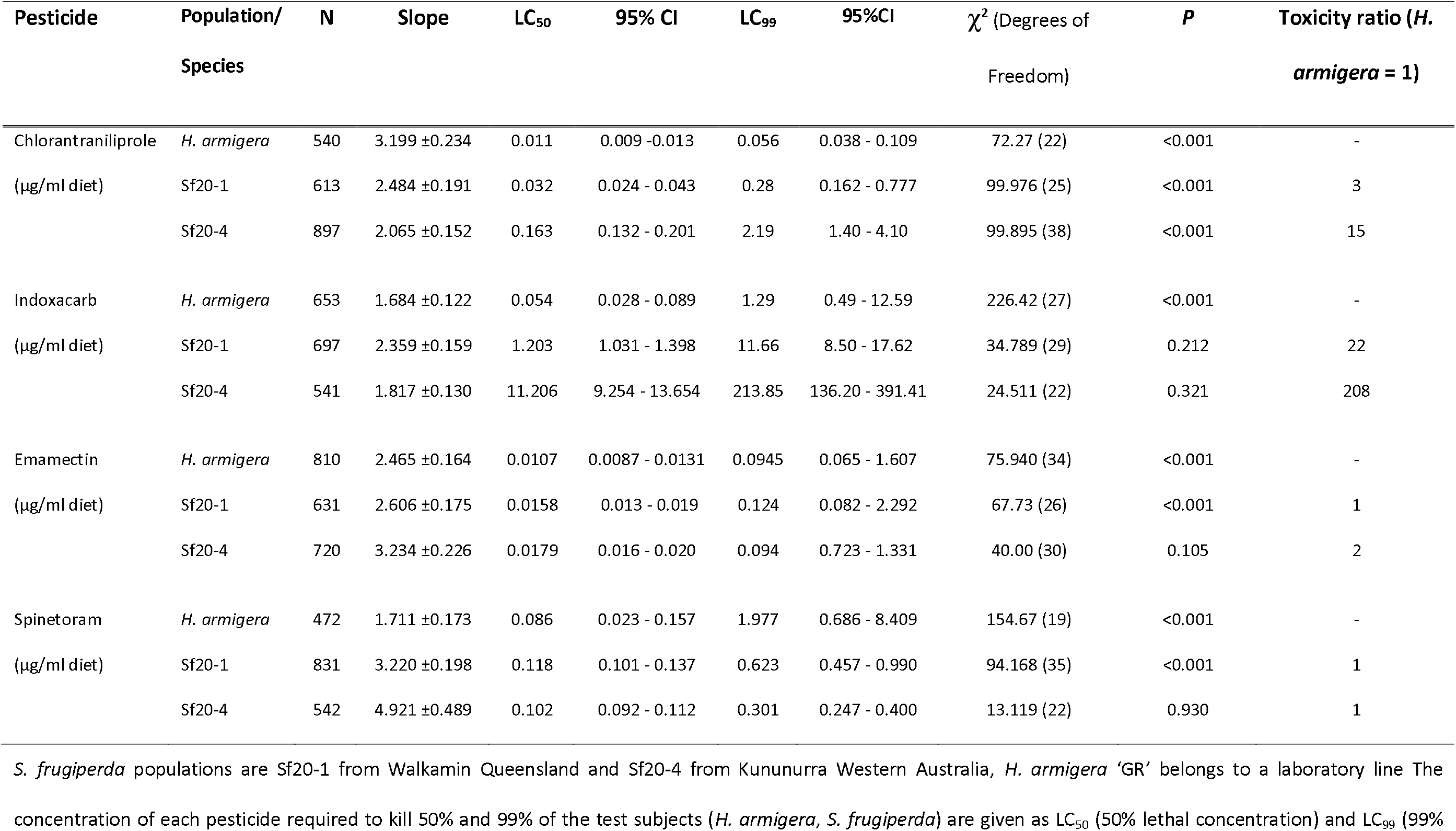
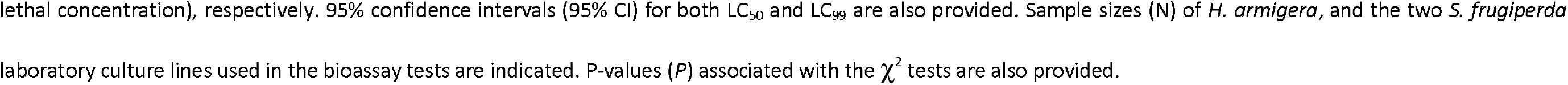
Summary bioassay data on *Spodoptera frugiperda* and *Helicoverpa armigera* involving diet incorporation of insecticides.

| Pesticide | Population/<br>Species | N | Slope | LC<sub>50</sub> | 95% CI | LC<sub>99</sub> | 95%CI | $\chi^2$ (Degrees of<br>Freedom) | P | Toxicity ratio (*H.*<br>*armigera* = 1) |
| --- | --- | --- | --- | --- | --- | --- | --- | --- | --- | --- |
| Chlorantraniliprole<br>($\mu\text{g/ml}$ diet) | *H. armigera* | 540 | 3.199 ± 0.234 | 0.011 | 0.009 - 0.013 | 0.056 | 0.038 - 0.109 | 72.27 (22) | <0.001 | - |
| | Sf20-1 | 613 | 2.484 ± 0.191 | 0.032 | 0.024 - 0.043 | 0.28 | 0.162 - 0.777 | 99.976 (25) | <0.001 | 3 |
| | Sf20-4 | 897 | 2.065 ± 0.152 | 0.163 | 0.132 - 0.201 | 2.19 | 1.40 - 4.10 | 99.895 (38) | <0.001 | 15 |
| Indoxacarb<br>($\mu\text{g/ml}$ diet) | *H. armigera* | 653 | 1.684 ± 0.122 | 0.054 | 0.028 - 0.089 | 1.29 | 0.49 - 12.59 | 226.42 (27) | <0.001 | - |
| | Sf20-1 | 697 | 2.359 ± 0.159 | 1.203 | 1.031 - 1.398 | 11.66 | 8.50 - 17.62 | 34.789 (29) | 0.212 | 22 |
| | Sf20-4 | 541 | 1.817 ± 0.130 | 11.206 | 9.254 - 13.654 | 213.85 | 136.20 - 391.41 | 24.511 (22) | 0.321 | 208 |
| Emamectin<br>($\mu\text{g/ml}$ diet) | *H. armigera* | 810 | 2.465 ± 0.164 | 0.0107 | 0.0087 - 0.0131 | 0.0945 | 0.065 - 1.607 | 75.940 (34) | <0.001 | - |
| | Sf20-1 | 631 | 2.606 ± 0.175 | 0.0158 | 0.013 - 0.019 | 0.124 | 0.082 - 2.292 | 67.73 (26) | <0.001 | 1 |
| | Sf20-4 | 720 | 3.234 ± 0.226 | 0.0179 | 0.016 - 0.020 | 0.094 | 0.723 - 1.331 | 40.00 (30) | 0.105 | 2 |
| Spinetoram<br>($\mu\text{g/ml}$ diet) | *H. armigera* | 472 | 1.711 ± 0.173 | 0.086 | 0.023 - 0.157 | 1.977 | 0.686 - 8.409 | 154.67 (19) | <0.001 | - |
| | Sf20-1 | 831 | 3.220 ± 0.198 | 0.118 | 0.101 - 0.137 | 0.623 | 0.457 - 0.990 | 94.168 (35) | <0.001 | 1 |
| | Sf20-4 | 542 | 4.921 ± 0.489 | 0.102 | 0.092 - 0.112 | 0.301 | 0.247 - 0.400 | 13.119 (22) | 0.930 | 1 |

*S. frugiperda* populations are Sf20-1 from Walkamin Queensland and Sf20-4 from Kununurra Western Australia, *H. armigera* 'GR' belongs to a laboratory line The concentration of each pesticide required to kill 50% and 99% of the test subjects (*H. armigera*, *S. frugiperda*) are given as LC<sub>50</sub> (50% lethal concentration) and LC<sub>99</sub> (99%
lethal concentration), respectively. 95% confidence intervals (95% CI) for both LC<sub>50</sub> and LC<sub>99</sub> are also provided. Sample sizes (N) of *H. armigera*, and the two *S. frugiperda* laboratory culture lines used in the bioassay tests are indicated. P-values (*P*) associated with the $\chi^2$ tests are also provided.

The dose response and LC_50_ for chlorantraniliprole show similar sensitivity for *H. armigera* and *S. frugiperda* Sf20-1, with the latter showing a 3x difference in the LC_50_. Interestingly, the *S. frugiperda* Sf20-4 exhibited a 15x reduction in sensitivity compared to *H. armigera* (Fig. S10; Table 5), and a 5x response variation to chlorantraniliprole when compared to the Sf20-1 line.

The dose response and LC_50_ data for emamectin benzoate and spinetoram (Figs. S11, S12; Table 5) suggest that *S. frugiperda* Sf20-1 and Sf20-4 are not significantly different in their level of sensitivity compared to *H. armigera*.

### Resistance alleles by whole genome sequencing

The loci examined for potential resistance alleles were present in the data set with the appropriate level of coverage to accurately call the genotype. Resistance alleles to carbamate/organophosphates (ACE-1) were the only resistance alleles identified in the Kununnura, Strathmore, South Korea, Papua New Guinea, and Peru populations. No resistance alleles associated with target site mutations were detected for pyrethroid (VGSC) or for diamide insecticides (RyR), although there are likely other genes (e.g., detoxification genes) that confer resistance to synthetic pyrethroid in these laboratory lines of *S. frugiperda* (Bird et al. 2022). Resistance allele profile differences between the two Australian populations were evident between the A201S and the F290V amino acid substitutions. Similar ACE-1 resistance allele profiles were detected between Strathmore and South Korean populations, while the Papua New Guinea population shared ACE-1 allele profiles with Strathmore population (A201S) and with Kununnura (F290V). The Peruvian *S. frugiperda* population was the only population that had heterozygous individuals with resistance allele for G227A amino acid substitution (Table 6).

**Table 6:** ACE-1 locus of *Spodoptera frugiperda* individuals from Australia, South Korea, Papua New Guinea, and Peru characterised via whole genome.

| Population | N | ACE-1 (A201S) |  |  | ACE-1 (G227A) |  |  | ACE-1 (F290V) |  |  |
| --- | --- | --- | --- | --- | --- | --- | --- | --- | --- | --- |
|  |  | S/S | S/R | R/R | S/S | S/R | R/R | S/S | S/R | R/R |
| Kununnura | 9 | 9 | 0 | 0 | 9 | 0 | 0 | 2 | 4 | 3 |
| Strathmore | 30 | 26 | 4 | 0 | 30 | 0 | 0 | 0 | 0 | 30 |
| South Korea | 12 | 11 | 1 | 0 | 12 | 0 | 0 | 0 | 0 | 12 |
| Papua New Guinea | 17 | 15 | 2 | 0 | 17 | 0 | 0 | 1 | 12 | 4 |
| Peru | 16 | 4 | 12 | 0 | 10 | 6 | 0 | 2 | 14 | 0 |

Number of *Spodoptera frugiperda* individuals (N) from populations from Australia (Kununnura SF20-4; Strathmore SF20-1), South Korea, Papua New Guinea, and Peru with homozygous susceptible (S/S), heterozygous (S/R) and homozygous resistance (R/R) profiles characterised via whole genome sequence data of the ACE-1 locus involving the A2001S, G227A, and F290V amino acid substitutions.

Resistance allele characterisation by sequencing approaches from this study and published studies for the ACE-1 gene is summarised in (Fig. 1, Table S2 and references therein). The most common resistance allele detected in the invasive and native population (456 individuals examined in total) was the F290V mutation with 66 homozygous and 222 heterozygous resistant genotypes detected. This mutation is a T to G single nucleotide polymorphism (SNP) that changes the codon encoding the amino acid from TTT to GTT leading to a phenylalanine (F) to valine (V) change in the protein sequence encoded by the ACE-1 gene. This mutation was present at all locations and evenly distributed between the invasive and native populations. In Australia, of the sequenced individuals (n = 146), 24.7% (n = 36) were heterozygous, and 8% (n = 12) were homozygous for the resistance allele. Heterozygous and homozygous individuals were found in populations from all four states (i.e., WA, NT, Queensland, and NSW) as well as from the single individual from Erub Island, suggesting it is common across Australia.

**Fig. 1:**
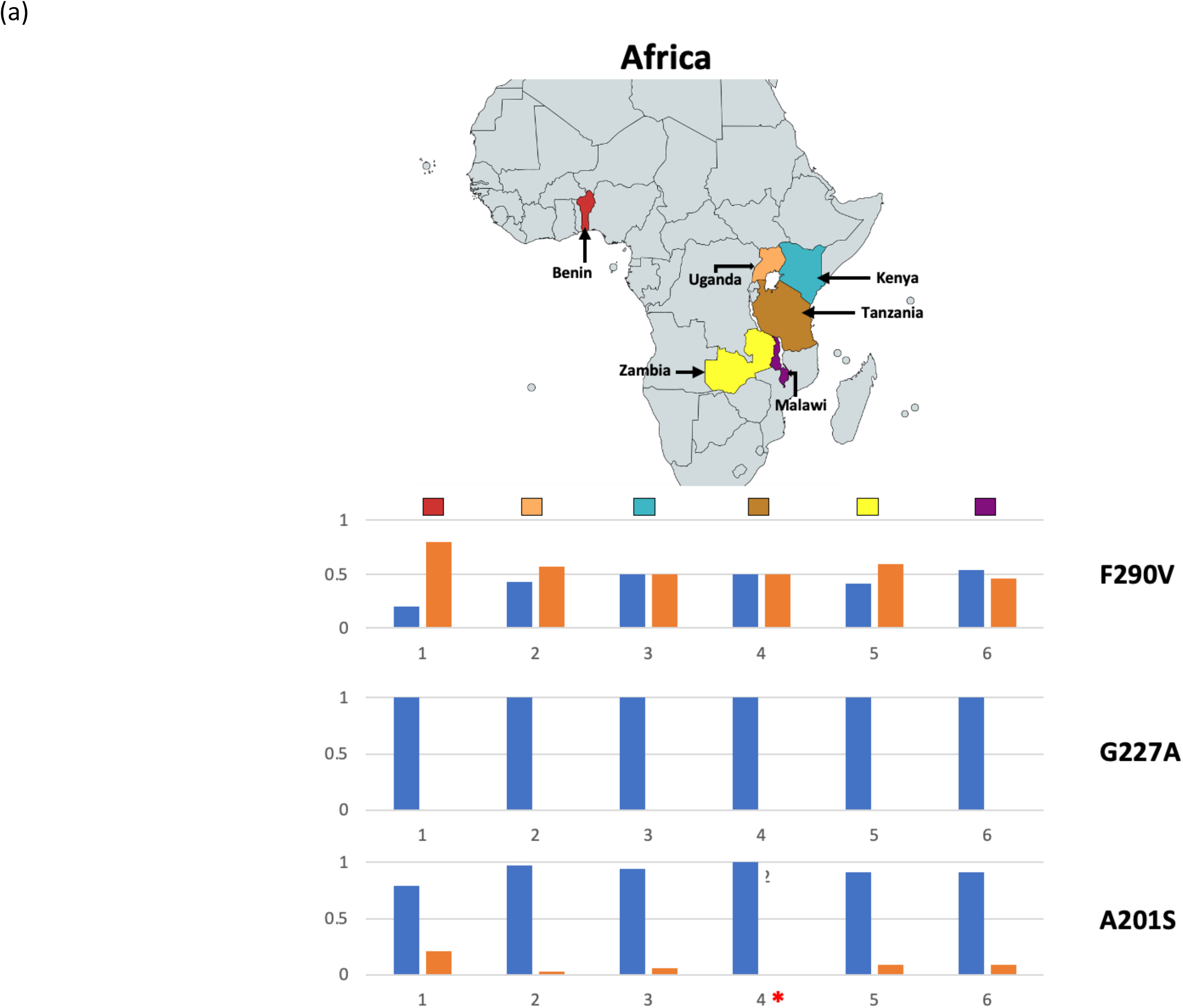
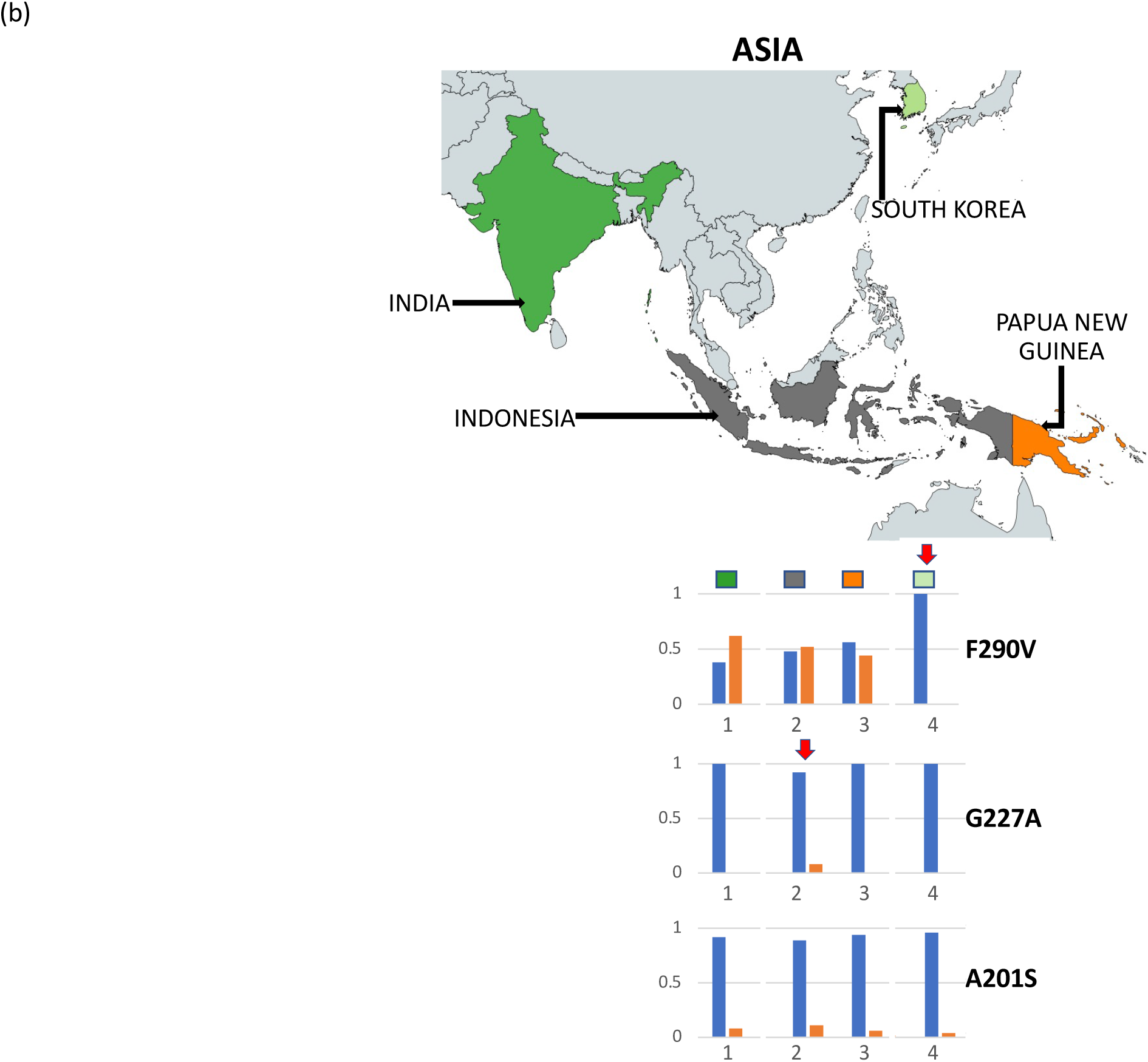
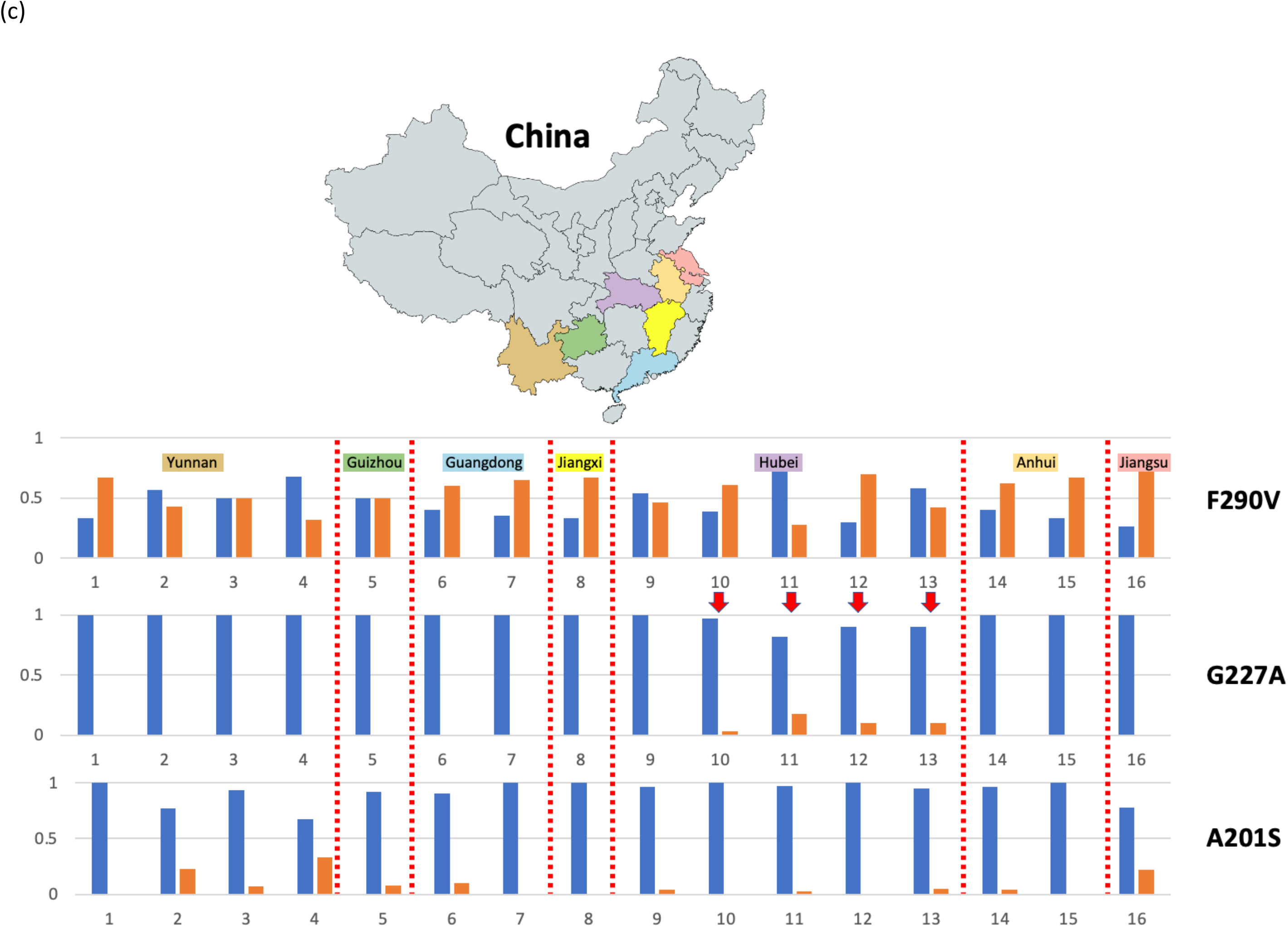
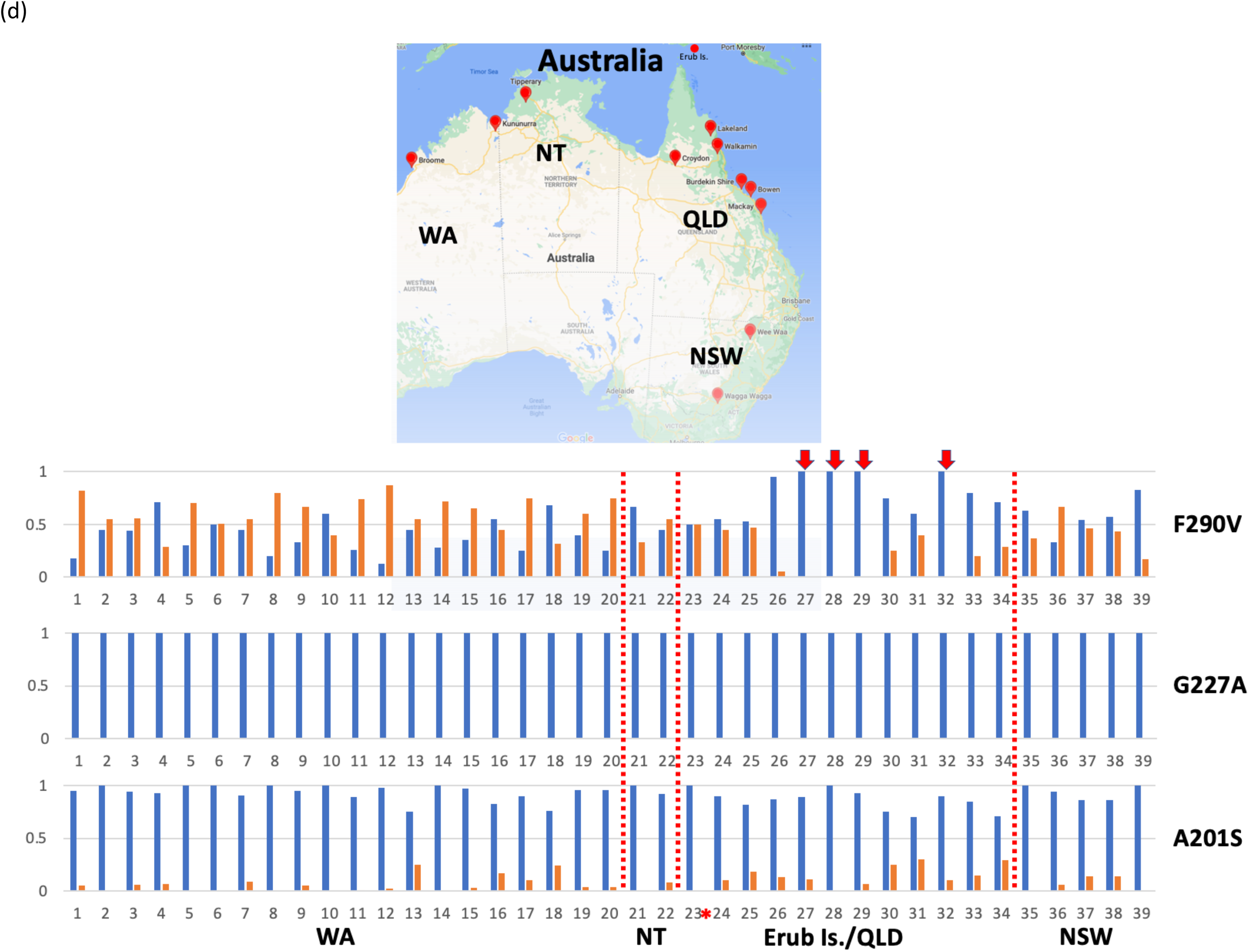
Summary of Acytylcholinesterase (ACE-1) susceptible and resistance allele frequencies in invasive range *Spodoptera frugiperda* populations from: (a) six African countries (i.e., Benin, Uganda, Kenya, Tanzania, Zambia, Malawi), (b) Asia (i.e., India, Indonesia, Papua New Guinea (PNG), South Korea), (c) China, and (d) Australia. A total of 1,177 individuals representing 75 populations in Table S2 were used to compile the data below. Population identity is as provided in Table S2 that combined data from this study (Australia SF20-1 and SF20-4 (generation 0 representing field-collected individuals), PNG, South Korea) and from published studies (Boaventura et al. 2020a, Guan et al. 2020, Zhang et al. 2020, Zhao et al. 2020, Nguyen et al. 2021, Tay et al. 2021, Yainna et al. 2021, Rane et al. 2022). Susceptible and resistant alleles from the three previously reported loci (i.e., F290V, G227A, A201S) from the ACE-1 gene provided evidence to support multiple independent introductions across the invasive *S. frugiperda* populations, such as in (b) Indonesia (#2; G227A) and South Korea (#4; F290V), and in (c) China (Hubei province (#10-13); G227A) as indicated by the red arrows. In (d) Australia, newly established *S. frugiperda* populations between Queensland (e.g., #27, #28 (Walkamin), #29 (Strathmore), #32 (Burdekin), and Western Australia (i.e., #2 (Kununurra)/Northern Territory (e.g., #21 (Bluey’s Farm)) suggested this likely involved multiple introductions from diverse populations from neighbouring countries and likely arrived via separate pathways and entry points (see also Rane et al. 2022).

The second most common allele detected was A201S (456 individuals examined) with 90 heterozygous individuals detected. This mutation is a C to A SNP which changes the codon from GCG to TCG leading to an alanine (A) to serine (S) amino acid change. While less common, this mutation also appears to be in both the invasive and native range with no obvious pattern. In Australia, of the sequenced individuals (n = 146), 15.8% (n = 23) were heterozygous, and 0 were homozygous for the resistance allele. Heterozygous individuals were found in WA and Queensland, also suggesting it is common across Australia.

The G227A mutation was the least common (from 456 individuals) with 20 heterozygous individuals and 2 homozygous individuals detected. This mutation is a G to a C SNP which alters the codon from GGA to CGA encoding a glycine (G) to alanine (A) amino acid change. Interestingly G227A was only present in individuals from the native range (Brazil, USA, Puerto Rico, Peru) but absent in individuals from across the invasive range that were surveyed in this study and from the related studies of Tay et al. (2021, 2022a). When compiling allele frequencies for both the VGSC and ACE-1 resistant genes from this study and from published whole genome sequencing and targeted PCR/Sanger sequencing data (Fig. 1; see also Table S1), the rare G227A resistance allele was present only in the Indonesian (Boaventura et al. 2020a) and Hubei populations (Guo et al. 2020) but absent in African, Australian, and other Asian (e.g., Indian, South Korea) populations, including populations from six other Chinese provinces.

No target site mutation alleles predicted to cause resistance to pyrethroids or the group 28 pesticides were detected in this work. However, while the previously identified resistance alleles (Bolzan et al. 2019, Boaventura et al. 2020b) were not detected, considerable variation was present in the RyR gene at the potential resistance loci. This should be further investigated in conjunction with bioassays to establish whether any of the variants could contribute to resistance.

### ABCC2 resistance alleles in Australia, Papua New Guinea, South Korea, and Peru populations

None of the known and validated Cry1 resistance ABCC2 mutations were identified in the individuals sequenced for this work however, some variation from the reference was observed (Fig. S13). Numerous synonymous mutations and a total of 31 non-synonymous mutations were observed in the coding sequence of the ABCC2 gene in all sequenced individuals. Several of these non-synonymous mutations were associated with a common deletion and insertion motif in the first exon where an 11bp deletion and a 2 bp insertion maintain the reading frame but replace and change several amino acids. Most of the other mutations are the result of one (i.e., single nucleotide variant; SNV) or two (i.e., multiple nucleotide variant; MNV) nucleotide changes. Several of these mutations are found in other assemblies of *S. frugiperda* and likely reflect natural variation. An alternative explanation for the variation is that they are associated with the c-strain which is thought to make up at least some of the genome of the invasive populations. Only one mutation, a 2 bp deletion in one individual collected from the Burdekin in Queensland was identified that might cause a frame shift mutation (see Supplemental Figs. S14 and S15) as has been identified in Bt resistant individuals in other studies. It was present as a heterozygote in the individual and has not been experimentally validated.

## Discussion

In this study, we showed insecticide and Bt response differences in two of the first reported *S. frugiperda* populations in Queensland (i.e., SF20-1) and Western Australia (Sf20-4) in Australia, with the Queensland (Sf20-1) population being less tolerant to various insecticides such as methomyl, chlorantraniliprole, and indoxacarb compared to the Western Australian (Sf20-4) population. On the other hand, the Queensland population exhibited at least a two-fold higher tolerance to Cry1Ac, Cry2Ab, Cry1F and Vip3A Bt toxins than the WA population. Characterisation of the ABCC2 resistance gene identified one *S. frugiperda* individual from Queensland as potentially being heterozygous with a 2 bp deletion that could underpin Cry1F resistance, although confirmation of the detected mutation and of the resistance phenotype is required. This could be accomplished via PCR and Sanger sequencing (e.g., Guan et al. 2020) as well as via the CRISPR/Cas9 gene editing approach (e.g., see Wang et al. 2017). The response differences to chemical insecticides and Bt toxins of the two studied invasive *S. frugiperda* populations in Australia and their ACE-1 resistance allele profile differences (Table 6) may reflect the diverse genetic composition across the pest’s recent expanding range (Tay et al. 2022a, Schlum et al. 2021, Rane et al. 2022), and suggest that separate pathways were involved in establishment of these Queensland and Western Australian populations. This is contrary to the current postulation of a single introduction pathway for the arrival of *S. frugiperda* to Australia based on an assumption (Jing et al. 2021) or reverse trajectory simulation (Qi et al. 2021) and highlights the importance of harmonising simulation studies with genomic and phenotypic evidence.

With significant economic impacts on agriculture from *S. frugiperda* reported in over 80 countries (excluding the New World native range) from Africa, Middle East, Asia, Southeast Asia, and Oceania, its response to different insecticidal and Bt control agents is increasingly investigated at recently impacted localities (e.g., Worku and Ebabuye 2019, Deshmukh et al. 2020, Zhang et al. 2020, Kulye et al. 2021, Lv et al. 2021). However, the diverse bioassay methods (approach, larval stage, scoring criteria) used in these studies complicate meaningfully comparison of findings. Taking emamectin benzoate and indoxacarb as examples, Zhang et al. (2022) used a topical application bioassay on 3^rd^ instar larvae, Deshmukh et al. (2020) used leaf-dip bioassays on 2^nd^ instar larvae, while Hardke et al. (2011) used a diet-incorporation approach on 3^rd^ instar larvae. Responses have been expressed as LD_50_ (e.g., Deshmukh et al. 2020) or as EC_50_ (e.g., Kulye et al. 2021), further making it difficult to rigorously compare outcomes.

It is important to stress the challenge and difficulty to compare bioassay results between studies due to the different genetic background of the test samples (e.g., due to different number of individuals used to establish test populations), methods and approaches between research groups, and the general different rearing conditions of laboratory cultures that could contribute to varied response outcomes. A further challenge for invasive *S. frugiperda* management is the lack of base-line values representing susceptible responses in newly populated areas which makes it difficult to monitor changes through time in insecticide efficacy due to resistance. Knowledge of whether the introduction occurred once (e.g., single ‘invasive bridgehead effect’; Guillemaud et al. 2011) vs. multiple times (e.g., multiple ‘mass dispersal’; Wilson et al. 2009) and patterns of gene flow also impact the long-term monitoring of insecticide resistance evolution. Using native *S. frugiperda* colonies established in 2005 from cotton fields in Louisiana, USA, Wilson et al. (2009) undertook diet-incorporation bioassays (on 3^rd^ instar larvae), using insecticides including chlorantraniliprole, indoxacarb, and spinetoram (see Table 7). Yu (1991) undertook topical application bioassays against methomyl (on 4^th^ instar larvae) which included comparison with a susceptible population free from insecticide exposure since 1975, and a resistant population collected from a maize field in Gainesville, Florida (see Table 8). In comparing insecticide responses in Indian invasive *S. frugiperda* populations with diet incorporation assays (on 3^rd^ instar larvae), Yu (1991) also included a native susceptible *S, frugiperda* population from Brazil to assist with interpreting changes to insecticide responses (including chlorantraniliprole and spinetoram) at spatial and temporal scales.

**Table 7:**
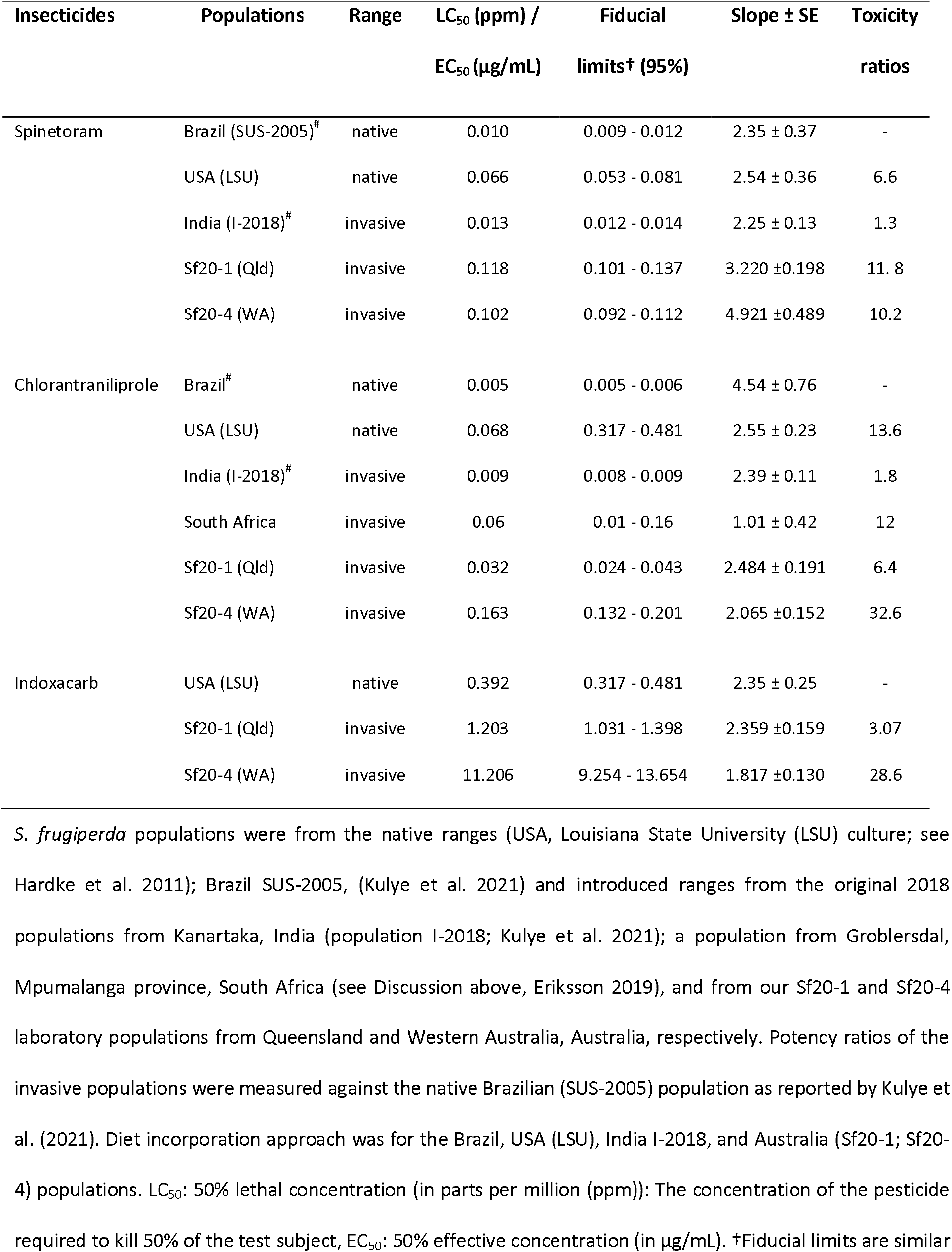
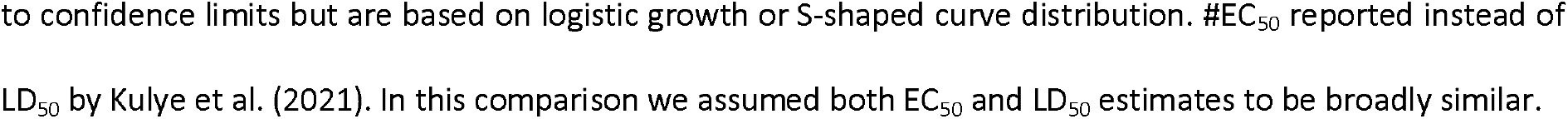
Comparisons of bioassay results for selected insecticides fed to laboratory populations of *Spodoptera frugiperda* via diet incorporation approaches.

| Insecticides | Populations | Range | LC<sub>50</sub> (ppm) /<br>EC<sub>50</sub> (µg/mL) | Fiducial<br>limits† (95%) | Slope ± SE | Toxicity<br>ratios |
| --- | --- | --- | --- | --- | --- | --- |
| Spinetoram | Brazil (SUS-2005)<sup>#</sup> | native | 0.010 | 0.009 - 0.012 | 2.35 ± 0.37 | - |
|  | USA (LSU) | native | 0.066 | 0.053 - 0.081 | 2.54 ± 0.36 | 6.6 |
|  | India (I-2018)<sup>#</sup> | invasive | 0.013 | 0.012 - 0.014 | 2.25 ± 0.13 | 1.3 |
|  | Sf20-1 (Qld) | invasive | 0.118 | 0.101 - 0.137 | 3.220 ± 0.198 | 11.8 |
|  | Sf20-4 (WA) | invasive | 0.102 | 0.092 - 0.112 | 4.921 ± 0.489 | 10.2 |
| Chlorantraniliprole | Brazil<sup>#</sup> | native | 0.005 | 0.005 - 0.006 | 4.54 ± 0.76 | - |
|  | USA (LSU) | native | 0.068 | 0.317 - 0.481 | 2.55 ± 0.23 | 13.6 |
|  | India (I-2018)<sup>#</sup> | invasive | 0.009 | 0.008 - 0.009 | 2.39 ± 0.11 | 1.8 |
|  | South Africa | invasive | 0.06 | 0.01 - 0.16 | 1.01 ± 0.42 | 12 |
|  | Sf20-1 (Qld) | invasive | 0.032 | 0.024 - 0.043 | 2.484 ± 0.191 | 6.4 |
|  | Sf20-4 (WA) | invasive | 0.163 | 0.132 - 0.201 | 2.065 ± 0.152 | 32.6 |
| Indoxacarb | USA (LSU) | native | 0.392 | 0.317 - 0.481 | 2.35 ± 0.25 | - |
|  | Sf20-1 (Qld) | invasive | 1.203 | 1.031 - 1.398 | 2.359 ± 0.159 | 3.07 |
|  | Sf20-4 (WA) | invasive | 11.206 | 9.254 - 13.654 | 1.817 ± 0.130 | 28.6 |

*S. frugiperda* populations were from the native ranges (USA, Louisiana State University (LSU) culture; see Hardke et al. 2011); Brazil SUS-2005, (Kulye et al. 2021) and introduced ranges from the original 2018 populations from Kanartaka, India (population I-2018; Kulye et al. 2021); a population from Groblersdal, Mpumalanga province, South Africa (see Discussion above, Eriksson 2019), and from our Sf20-1 and Sf20-4 laboratory populations from Queensland and Western Australia, Australia, respectively. Potency ratios of the invasive populations were measured against the native Brazilian (SUS-2005) population as reported by Kulye et al. (2021). Diet incorporation approach was for the Brazil, USA (LSU), India I-2018, and Australia (Sf20-1; Sf20-4) populations. LC<sub>50</sub>: 50% lethal concentration (in parts per million (ppm)); The concentration of the pesticide required to kill 50% of the test subject, EC<sub>50</sub>: 50% effective concentration (in µg/mL). †Fiducial limits are similar
to confidence limits but are based on logistic growth or S-shaped curve distribution. #EC<sub>50</sub> reported instead of LD<sub>50</sub> by Kulye et al. (2021). In this comparison we assumed both EC<sub>50</sub> and LD<sub>50</sub> estimates to be broadly similar.

**Table 8:** Comparisons of topically applied methomyl bioassay on *S. frugiperda* between resistant or susceptible native populations from USA (FL) and invasive populations from South and Australia.

| Populations | LC<sub>50</sub> (ppm) | Fiducial limits (95%) | Slope ± SE | Toxicity ratios |
| --- | --- | --- | --- | --- |
| USA (FL susceptible) | 0.57 | 0.45 - 0.78 | 4.9 | - |
| South Africa | 62.98 | 17.22 – 52.12 | 11.14 ± 9.20 | 110.5 |
| Sf20-1 (Qld) | 0.254 | 0.177 - 0.356 | 1.064 ± 0.076 | 0.45 |
| Sf20-4 (WA) | 2.96 | 1.87 - 4.66 | 0.874 ± 0.07 | 5.2 |
| USA (FL resistance) | 8.23 | 4.86 - 11.6 | 2.9 | - |
| South Africa | 62.98 | 17.22 – 52.12 | 11.14 ± 9.20 | 7.65 |
| Sf20-1 (Qld) | 0.254 | 0.177 - 0.356 | 1.064 ± 0.076 | 0.031 |
| Sf20-4 (WA) | 2.96 | 1.87 - 4.66 | 0.874 ± 0.07 | 0.36 |

Comparisons of topically applied methomyl bioassay results on late 3<sup>rd</sup> and/or early 4<sup>th</sup> instar larvae of *S. frugiperda* between resistant or susceptible populations from Gainesville Florida, USA (FL) (Yu 1991; see Discussion above), and invasive populations from South Africa (Eriksson 2019; see Discussion above) and Queensland or Western Australia, Australia. LC<sub>50</sub> (50% lethal concentration): The concentration of each pesticide required to kill 50% of the test subject. Fiducial limits are similar to confidence limits but are based on logistic growth or S-shaped curve distribution.

In the absence of native *S. frugiperda* populations in Australia, and in addition to the information collected herein for *H. armigera* and *S. litura*, we used the information from these global studies on native and newly invaded populations to assist interpretation of the early data collected in this study on invasive *S. frugiperda* populations in Australia. These examples used broadly similar approaches, instars and scoring criteria to our study for bioassays with specific insecticides. Relative to the indoxacarb results from Hardke et al. (2011) on native *S. frugiperda* populations from the USA, our bioassay findings for the invasive Western Australian and Queensland populations suggest a 28- and 3-folds difference, respectively (Table 7). For spinetoram, resistance ratios of the two Australia populations (which were like each other) is around 10 and 1.5 times higher than for the native populations from Brazil and the USA respectively. In contrast the ratios for the invasive populations from India was 1.3 and 0.3 relative to the Brazil and USA native populations. The differences are even more pronounced for chlorantraniliprole where the Sf20-4 WA population exhibited ratios that were 32 and 2.4 times higher than both the native Brazilian and American populations, which are at least around 3-fold higher than for the invasive populations in South Africa and India (Table 7) and also for the invasive Indian population studied by Deshmukh et al. (2020) using a leaf dip assay (and hence not reported in Table 7).

Methomyl resistance alleles have been reported in invasive populations from China, Indonesia, Africa, and in this study. Comparisons between susceptible and resistant strains of native *S. frugiperda* populations from Florida USA (Yu 1991) and with the South African invasive population of *S. frugiperda* (Eriksson 2019) showed that the Queensland population as similar in its response as the susceptible Florida *S. frugiperda*, while the Western Australia population was around 5 times more tolerant. This contrasted with the South African population which was 110 times more tolerant than the Florida susceptible *S. frugiperda* strain, and around 8 times more tolerant than the resistant *S. frugiperda* strain from Florida (Table 8). Taken as a whole, similar bioassay results between native (i.e., Brazil, USA) and various invasive populations suggested potential significant genetic diversity in introduced populations. This concurs with population genomic and genetic analyses (Tay et al. 2022a, Zhang et al. 2020, Schlum et al. 2021, Rane et al. 2022, Jiang et al. 2022) that suggested multiple origins for the invasive African, Asian (Indian, Chinese), and Southeast Asian (e.g., Malaysia) *S. frugiperda* populations.

It is possible that the differential response between the two Australian *S. frugiperda* populations to insecticides and Bt toxins represents natural variation in the national population which has emerged from a single founding incursion. However, it is unlikely that the differential responses within Australian populations represents local selection pressures because the period between populations establishing and being collected for this study was presumed to be too short to enable this opportunity. It could be that separate incursions involving different source populations occurred in WA compared to the eastern states of Australia, as supported by genome-wide SNP marker population genomic studies (Rane et al. 2022). Different selection pressures on the global population which recently originated from multiple Asian, Southeast Asian, and African incursions (Tay et al. 2022a, Zhang et al. 2020, Schlum et al. 2021, Rane et al. 2022) may have driven different phenotypes which entered the expanding ranges tested in our study and others reported herein. However, Kulye et al. (2021) demonstrated in *S. frugiperda* populations collected in India during 2018, 2019, and 2020 that large response changes such as those observed in South African (e.g., methomyl; Eriksson 2019) and Western Australian (chlorantraniliprole, indoxacarb; this study) populations were unlikely considering the short time frame since the very recent arrival of *S. frugiperda* especially assuming a west-to-east spread (Goergen et al. 2016, Cock et al. 2017, Nagoshi et al. 2018, 2019b). Another explanation for the differential responses of the two Australian populations that we studied is that they have different fitness because of variation in genetic diversity which reflects the number of individuals founding the populations. However, if this was the case one might expect sensitivity levels to be consistently higher or lower to all of the agents that we tested but this was not the case.

Alien invasive agricultural pests are increasingly being shown to carry novel insecticide resistance genes (e.g., Anderson et al. 2018, Walsh et al. 2018, Tay and Gordon 2019). This includes *S. frugiperda* in which invasive populations have been confirmed via whole genome sequence analyses (e.g., Guan et al. 2020, Zhang et al. 2020, Yainna et al. 2021) and molecular characterisation to harbour selected resistance genes (Boaventura et al. 2020a, 2020b; Zhao et al. 2020). Our review of reported ACE-1 and VGSC resistance allele frequency differences (Fig, 1; Table S2 and references therein) suggests that the invasive *S. frugiperda* populations within China (Guo et al. 2020), Indonesia (Boaventura et al. 2020a), Queensland, Australia (Tay et al. 2021, Rane et al. 2022), and South Korea (this study) were genetically diverse and likely originated from different native populations (Fig, 1). This further supports the perceived rapid spread of *S. frugiperda* across Africa, Asia, and Oceania as likely to also involve multiple independent introduction events.

Given the insecticide resistance allele frequency differences in *S. frugiperda* (e.g., Boaventura et al. 2020a, Guan et al. 2020, Lv et al. 2021, Yainna et al. 2021) and variation between the two populations sampled herein in bioassay responses to some of the approved insecticides for broadacre cropping, effort in Australia and indeed, for other invasive regions where possible, should now be directed to establishing baseline susceptibilities against key chemistries for multiple populations across geographies. For instance, pyrethroid resistance is common in the field in *H. armigera* (Walsh et al. 2018), and our comparisons with *H. armigera* and *S. litura* suggest that pyrethroids like cypermethrin are unlikely to provide good control in Australia against *S. frugiperda.* It also indicates some heterogeneity in the response to pyrethroids in the first invasive *S. frugiperda* populations in Australia. The large discrepancies between WA and Qld populations for methomyl and indoxacarb also require study of further populations before drawing firm conclusions. This information will be critical for on-going monitoring of resistance allele frequencies to key chemistries and determining their field application efficiencies, among geographies which is required to inform resistance management plans. It will also be important to understand gene flow patterns between different populations. Early detection of potential future introductions into Australia of novel resistance genes/alleles should also be a priority; for example, the VGSC L1014 resistance allele and the ACE G227A resistance allele from Southeast Asia (Boaventura et al. 2020a) and China (Guo et al. 2020), ryanodine receptor (RyR) resistance alleles from Brazil and China (Bolzan et al. 2019, Boaventura et al. 2020b, Lv et al. 2021) and ABCC2 resistance alleles from the Americas (Banerjee et al. 2017, Flagel et al. 2018, Guan et al. 2020, Yainna et al. 2021).

Being alert to new *S. frugiperda* incursions carrying novel resistance genes is relevant also for the Bt toxins. Our bioassay findings suggest Australia’s *S. frugiperda* populations likely do not carry Cry1F resistance alleles in the homozygous state known to exist in native range *S. frugiperda* populations (e.g., Banerjee et al. 2017, Flagel et al. 2018, Guan et al. 2020, Yainna et al. 2021), however the 2 bp deletion identified via whole genome sequencing in a single heterozygous individual from Burdekin (Queensland) will require further confirmation. Field-selected lines of VIP3Aa20 resistant *S. frugiperda* have been reported in Brazil and the USA where they occur as a recessive trait (Yang et al. 2013, Bernardi et al. 2015, 2016; Yang et al. 2018, 2019). The candidate resistance gene(s) and associated mutation(s) underpinning this resistance are yet to be identified, therefore phenotypic bioassays are required to detect resistance to VIP3A. Using what was essentially an F_0_ screen of material we did not detect resistance to VIP3A in our field-derived laboratory-maintained S. *frugiperda* colonies. However, bioassays involving F_2_-crosses (Andow and Alstad 1998) would be needed to further confirm the status of VIP3A resistance allele in both populations. The F_2_-crosses approach should be a valuable tool in protecting the Australian cotton industry against *S. frugiperda*, given the > 90% up-take of Bollgard III cotton containing Cry1Ac, Cry2Ab and Vip3A proteins. To protect agriculture in Australia (and elsewhere) from an accidental introduction of VIP3A resistant populations from the Americas (e.g., from Brazil, Bernardi et al. 2016) into global invasive populations, national (e.g., pre-border) and industry biosecurity preparedness strategies must be co-ordinated to prevent and to increase the chances of early detection of such novel introductions of new traits.

Novel introductions leading to unique population structure has been reported for western African *S. frugiperda* (Nagoshi et al. 2022), in populations from China (Jiang et al. 2022), in Africa (e.g., Benin vs. Malawi; Tay et al. 2022a), and in, e.g., Australia, Malaysia, and Myanmar vs. China populations (Rane et al. 2022), suggesting that the widely anticipated long distance migration of *S. frugiperda* especially in the invasive range could likely be less and may be impacted by localised ecological and climatic determinants (Tay et al. 2022b). Factors impacting the founding for the two Australian populations (e.g., founding number (i.e., involving one/few individuals vs. many); gene flow dynamics at spatial and temporal scales, and frequencies and impact from potential novel and on-going introductions) used in our bioassay studies is at present unknown. Bioassays testing of a larger population pooled from several sites or several individual populations from multiple sites would be especially relevant in a pest capable of high dispersal across the landscape, however the spread of *S. frugiperda* in the invasive range is increasingly being recognised as may not be as rapid and widespread as originally believed (e.g., Naghoshi et al. 2022; Rane et al. 2022; Tay et al. 2022a; Jiang et al. 2022). As such, we advocate cautionary approaches to avoid premature assumptions that extensive population admixture via gene flow in Australia landscape would lead to homogenised populations and therefore the approach to combine multiple distantly sampled populations in bioassay studies (e.g., Bird et al. 2022).

Finally, to enable meaningful comparisons of bioassay findings from across the *S. frugiperda* invasive range, there is a need to globally standardise approaches for testing of insecticides and Bt toxins. Localised novel resistance traits in populations of *S. frugiperda* would likely serve as a new resistance management challenge to other neighbouring regions (Kalyebi 2020), and movements of *S. frugiperda* in the new invasive ranges could lead to as yet unknown and complex gene flow patterns that could significantly hinder the development of suitable pest and resistance management strategies for this global pest complex.

## Acknowledgements

The *Spodoptera frugiperda* larvae used to establish the SF20-1 colony were provided by Prof. Myron Zalucki (University of Queensland). Mr Jack Daniel (Northern Australian Crop Research Alliance Pty Ltd) provided the *S. frugiperda* larvae used to establish SF20-4. Material from an established colony of *S. litura* was provided by Dr Ian Newton and Dr Melina Miles (QDAF). Some of the results reported herein were collected for a project led by CSIRO as a co-investment with the Grains Research and Development Corporation, the Australian Centre for International Agricultural Research, the Cotton Research and Development Corporation, FMC Australasia and Corteva Agriscience. GRDC manages the project on behalf of the co-investors.

