## Supplementary material for "Resistance bioassays and allele characterisation inform analysis of *Spodoptera frugiperda* (Lepidoptera: Noctuidae) introduction pathways in Asia and Australia": Table S1

**Supplementary Table S1**

**Resistance bioassays and allele characterisation inform analysis of *Spodoptera frugiperda* (Lepidoptera: Noctuidae) introduction pathways in Asia and Australia**

W. T. Tay^1,3^, R. V. Rane^2,3^, W. James^1^, K. H. J. Gordon^1^, S. Downes^4^, J. Kim^5^, L. Kuniata^6^, T. K. Walsh TK^1,3^

1. CSIRO Black Mountain Laboratories, Clunies Ross Street, ACT 2601, Australia

2. CSIRO, 343 Royal Parade, Parkville, VIC 3052, Australia

3. Applied BioSciences, Macquarie University, Sydney NSW 2100, Australia

4. CSIRO Mc Master Laboratories, New England Highway, Armidale NSW 2350, Australia

4. College of Agriculture and Life Science, Kangwon National University, Republic of Korea

5. Ramu Agri Industries Ltd., PNG

**Table S1:** SNPs used to identify resistance alleles. ACE-1 = the acetylcholine esterase gene 1; VGSC = the voltage gated sodium channel; RyR = the ryanodine receptor; ABCC2 = ATP Binding Cassette Subfamily C Member 2. Contig name is based on original *Spodoptera frugiperda* R-strain genome assembly (Gouin et al. 2017) used for analysis.

| Gene | Predicted Locus | Contig Name:SNP location | Pesticide |
| --- | --- | --- | --- |
| ACE-1 | G227A | SFRU_RICE_003833:20445 | Organophosphate/carbamate |
| ACE-1 | F290V | SFRU_RICE_003833:20258 | Organophosphate/carbamate |
| ACE-1 | A201S | SFRU_RICE_003833:20524 | Organophosphate/carbamate |
| VGSC | L932F | SFRU_RICE_011280:20607 | Pyrethroid |
| VGSC | L932F | SFRU_RICE_011280:20608 | Pyrethroid |
| VGSC | L932F | SFRU_RICE_011280:20609 | Pyrethroid |
| VGSC | L1014F | SFRU_RICE_011280:22407 | Pyrethroid |
| VGSC | L1014F | SFRU_RICE_011280:22408 | Pyrethroid |
| VGSC | L1014F | SFRU_RICE_011280:22408 | Pyrethroid |
| VGSC | T929I | SFRU_RICE_011280:20593 | Pyrethroid |
| VGSC | T929I | SFRU_RICE_011280:20594 | Pyrethroid |
| VGSC | T929I | SFRU_RICE_011280:20595 | Pyrethroid |
| RyR | I4790M | SFRU_RICE_011282:4512 | Group 28 |
| RyR | I4790M | SFRU_RICE_011282:4513 | Group 28 |
| RyR | I4790M | SFRU_RICE_011282:4514 | Group 28 |
| RyR | G4946E | SFRU_RICE_011282:2277 | Group 28 |
| RyR | G4946E | SFRU_RICE_011282:2278 | Group 28 |
| RyR | G4946E | SFRU_RICE_011282:2279 | Group 28 |
| ABCC2 | P799K/R | SFRICE002622:20935..20934 | Cry1F |
| ABCC2 | GY del | SFRICE002622:20947..20952 | Cry1F |
| ABCC2 | G1087D | SFRICE002622:18125..18127 | Cry1F |
| ABCC2 | Indel fs 865 | SFRICE002622:20583..20588 | Cry1F |
| ABCC2 | Indel fs 741 | SFRICE002622:21204..21209 | Cry1F |
