## Supplementary material for "Resistance bioassays and allele characterisation inform analysis of *Spodoptera frugiperda* (Lepidoptera: Noctuidae) introduction pathways in Asia and Australia": Fig. S1

**Fig. S1:** Cry1Ac dose response curves using diet overlay for *Helicoverpa armigera* (blue), Sf20-1 *Spodoptera frugiperda,* Walkamin, QLD (red), Sf20-4 *Spodoptera frugiperda,* Kununurra, WA (Black), *Spodoptera litura,* Mareeba, QLD (green). Percent response = mortality.

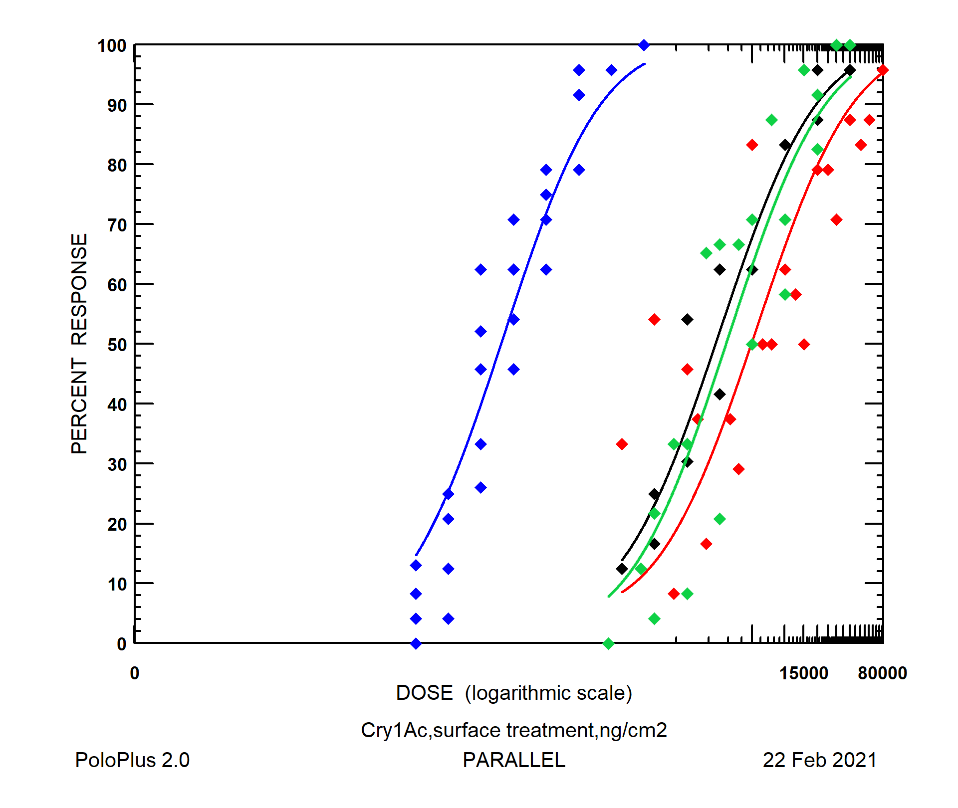
