## Supplementary material for "Resistance bioassays and allele characterisation inform analysis of *Spodoptera frugiperda* (Lepidoptera: Noctuidae) introduction pathways in Asia and Australia": Fig. S3

**Fig. S3:** Cry1F dose response curves using diet overlay for Sf20-1 *Spodoptera frugiperda*, Walkamin, QLD (red), Sf20-4 *Spodoptera frugiperda,* Kununurra, WA (Black), *Spodoptera litura,* Mareeba, QLD (green). Percent response = mortality

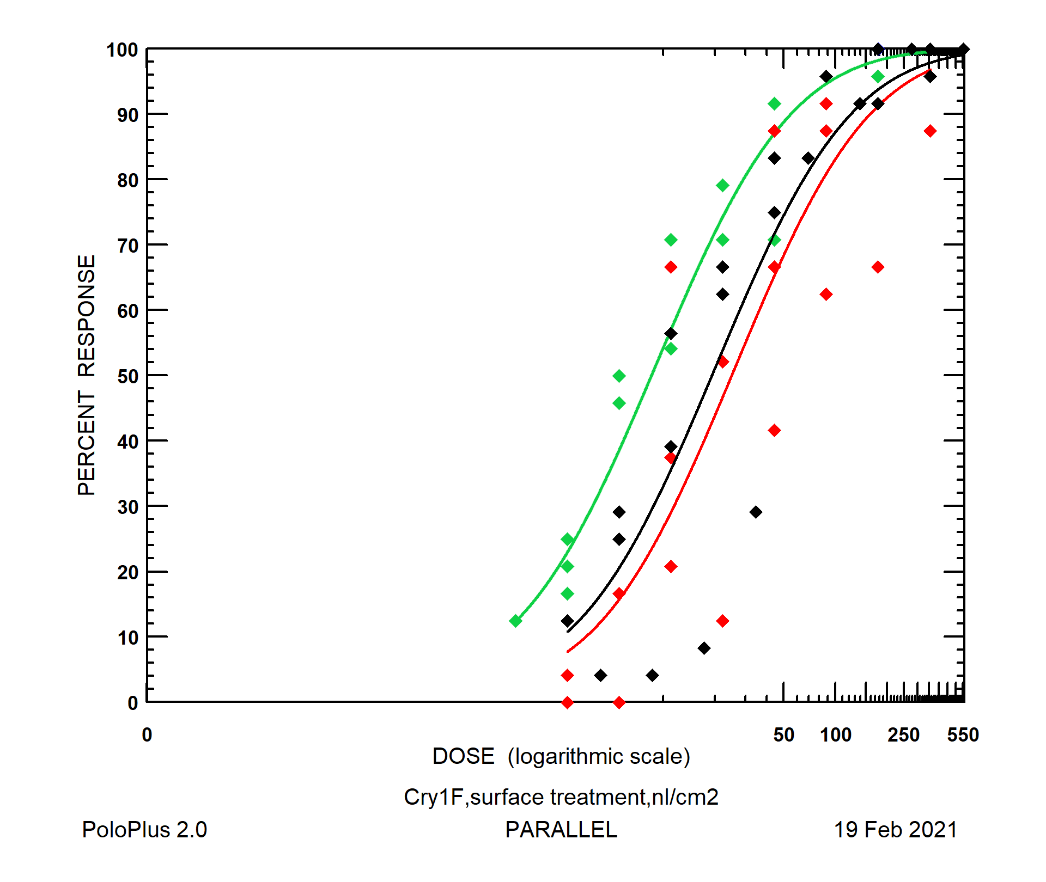
