## Supplementary material for "Resistance bioassays and allele characterisation inform analysis of *Spodoptera frugiperda* (Lepidoptera: Noctuidae) introduction pathways in Asia and Australia": Fig. S4

**Fig. S4:** VIP3a dose response curves using diet overlay for *Helicoverpa armigera* (blue), *Spodoptera frugiperda*, Walkamin, QLD (red), *Spodoptera frugiperda*, Kununurra, WA (Black), *Spodoptera litura*, Mareeba, QLD (green). Percent response = mortality.

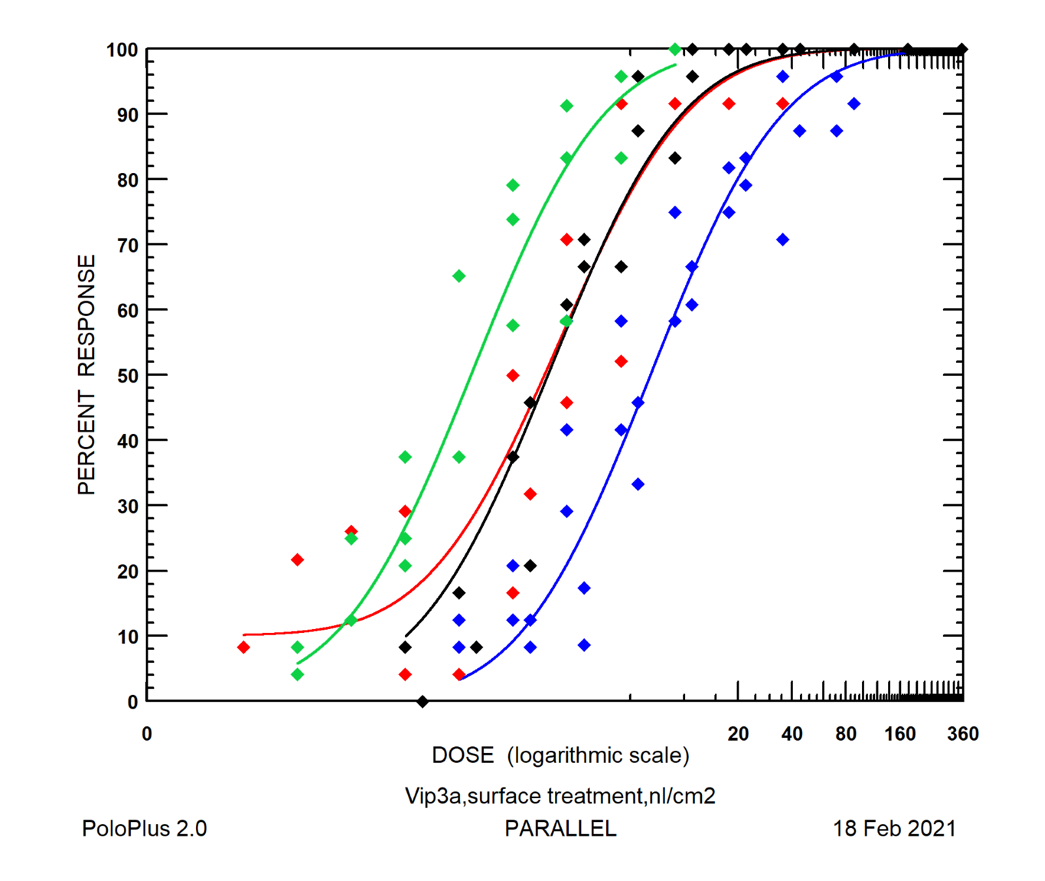
