## Supplementary material for "Resistance bioassays and allele characterisation inform analysis of *Spodoptera frugiperda* (Lepidoptera: Noctuidae) introduction pathways in Asia and Australia": Fig. S5

**Fig. S5:** Dipel dose response curves using diet overlay for *Helicoverpa armigera* (blue), Sf20-1 *Spodoptera frugiperda*, Walkamin, QLD (red), Sf20-4 *Spodoptera frugiperda*, Kununurra, WA (Black), *Spodoptera litura,* Mareeba, QLD (green). Percent response = mortality.

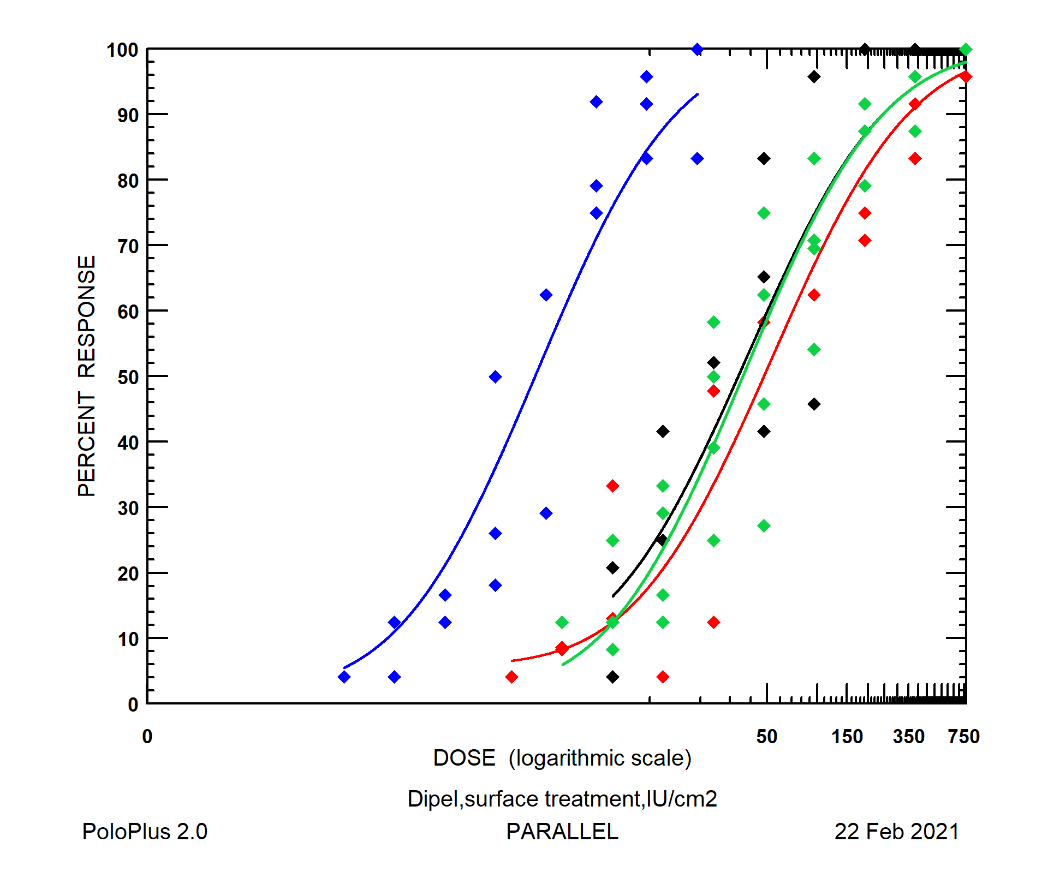
