## Supplementary material for "Resistance bioassays and allele characterisation inform analysis of *Spodoptera frugiperda* (Lepidoptera: Noctuidae) introduction pathways in Asia and Australia": Fig. S7

**Fig. S7:** Alpha-Cypermethrin dose response curves using diet incorporation for *Helicoverpa armigera* (blue), *Sf20-1 Spodoptera frugiperda, Walkamin, QLD (red), Sf20-4 Spodoptera frugiperda, Kununurra, WA (Black)*. Percent response = mortality.

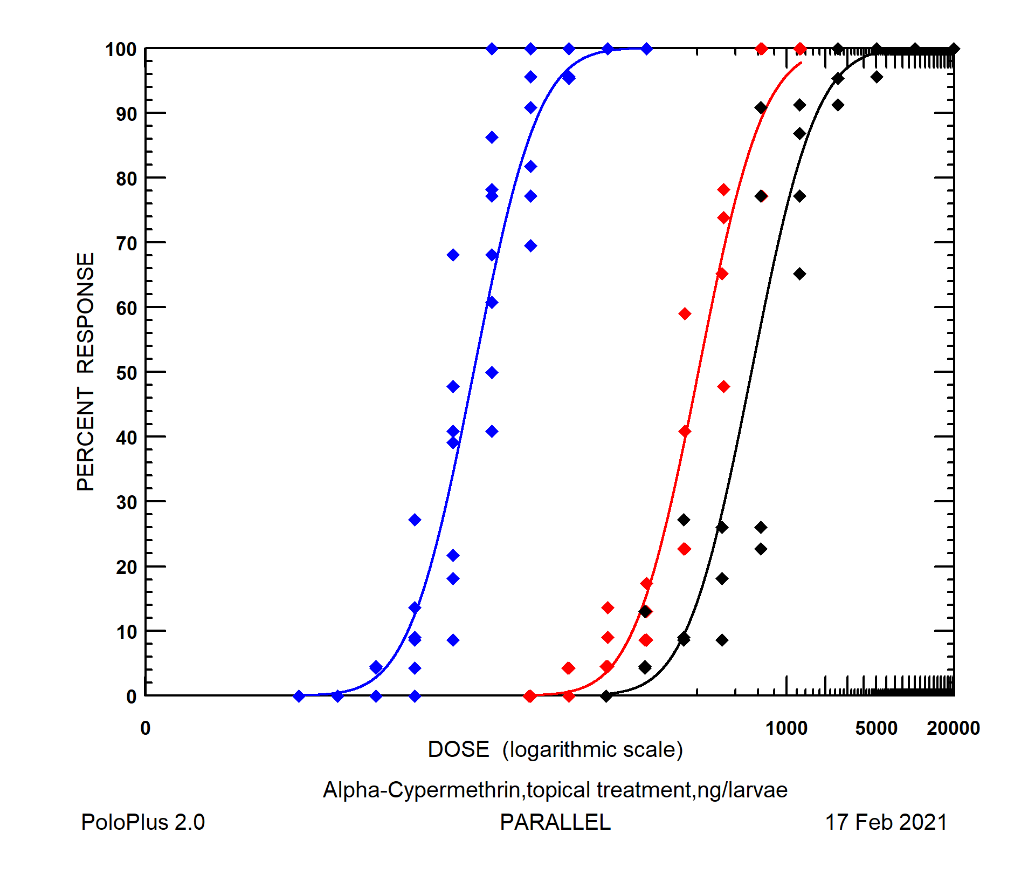
