## Supplementary material for "Resistance bioassays and allele characterisation inform analysis of *Spodoptera frugiperda* (Lepidoptera: Noctuidae) introduction pathways in Asia and Australia": Fig. S10

**Fig. S10:** Chlorantraniliprole dose response curves using diet incorporation for *Helicoverpa armigera* (blue), Sf20-1 *Spodoptera frugiperda*, Walkamin, QLD (red), Sf20-4 *Spodoptera frugiperda,* Kununurra, WA (Black). Percent response = mortality.

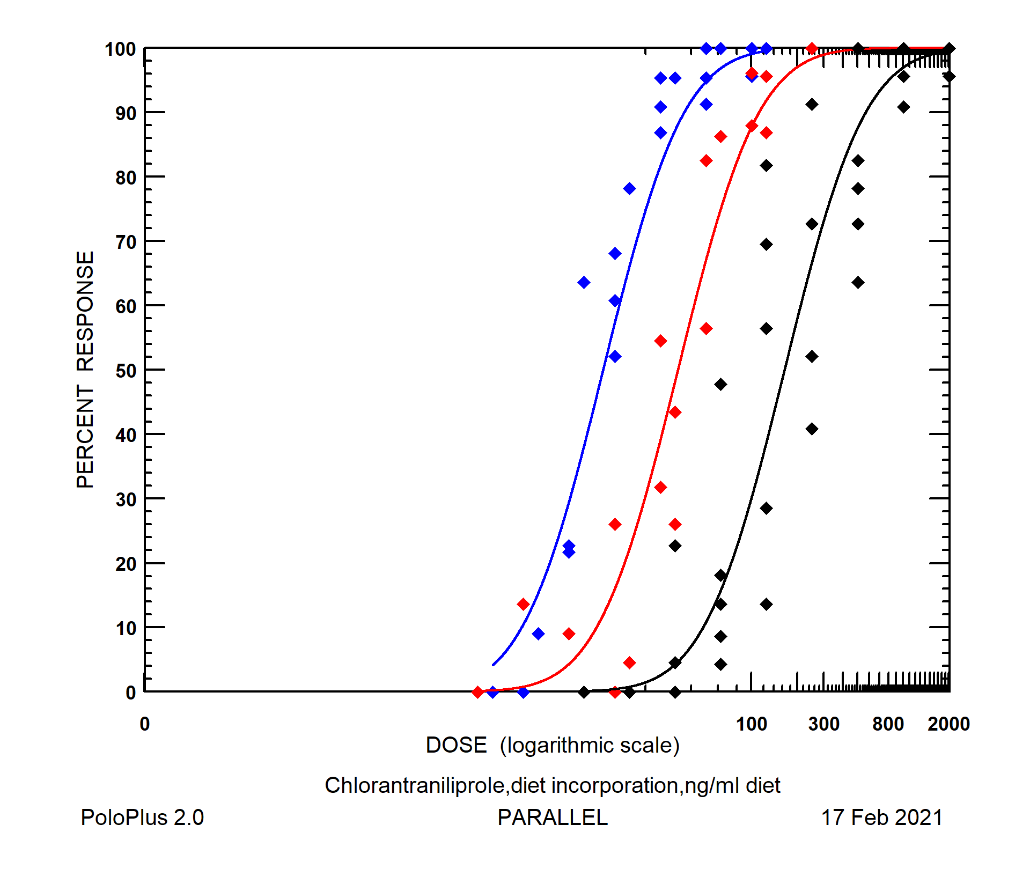
