## Supplementary material for "Resistance bioassays and allele characterisation inform analysis of *Spodoptera frugiperda* (Lepidoptera: Noctuidae) introduction pathways in Asia and Australia": Fig. S13

**Fig. S13:** The amino acid sequence of the ABCC2 gene from the R-strain *Spodoptera frugiperda* (SFRICE002622) (Gouin et al. 2017) aligned to the predicted amino acid sequence from an invasive genome (XP_035429361.1) (Xiao et al. 2020). In yellow are the differences between the two predicted seqs and on top are the amino acid changes observed in this dataset (i.e., Benin, Uganda, Kenya, Tanzania, Zambia, Malawi, India, Indonesia, Papua New Guinea, South Korea, China, Australia). In grey and # (indel) are the identified r alleles from Boaventura et al. (2020a) and Yainna et al. (2021). A number of novel substitutions and one potential disruptive mutation were observed. The potential Indel (in red) indicates the location of a potential frameshift mutation in an individual from QLD however, this was only observed in a single individual as a heterozygote with low coverage and further genetic and phenotypic validation would be required.

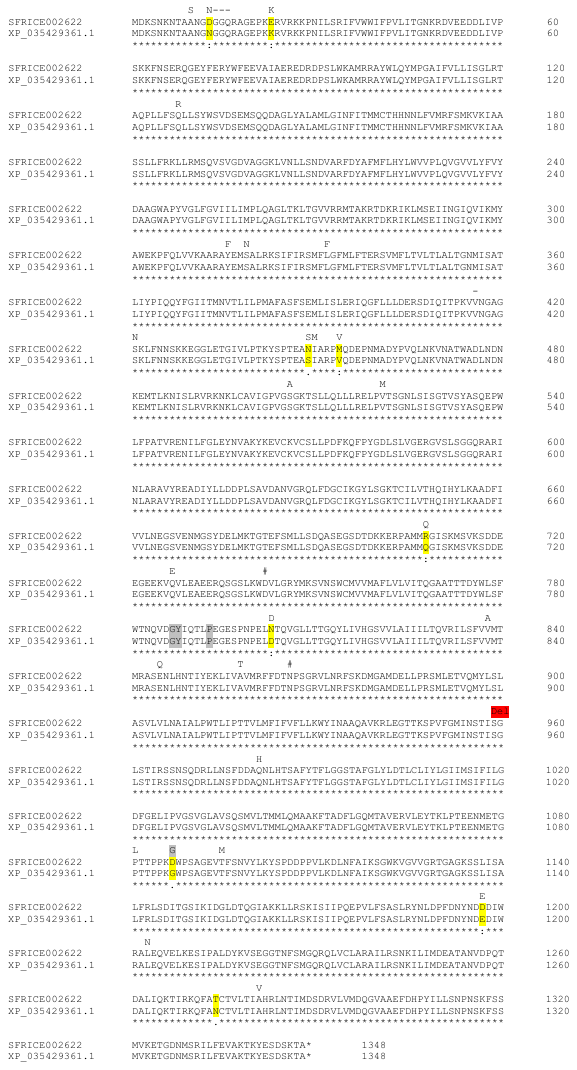
