## Supplementary material for "Resistance bioassays and allele characterisation inform analysis of *Spodoptera frugiperda* (Lepidoptera: Noctuidae) introduction pathways in Asia and Australia": Fig. S14

**Fig. S14:** Alignment of the coding sequence of the potential ABCC2 frameshift mutation in *Spodoptera frugiperda* from Australia. The two base pair deletion is highlighted in the wild type (WT) sequence with the corresponding amino acid change and eventual frameshift visible in the resistance allele (Res).

```
WT  cga ttg gaa gga aca act aag agt cca gtg ttt gga atg atc aac tct act atc tca gga 2880
WT   R   L   E   G   T   T   K   S   P   V   F   G   M   I   N   S   T   I   S   G  960
Res cga ttg gaa gga aca act aag agt cca gtg ttt gga atg atc aac tct act atc tgg act 2878
Res  R   L   E   G   T   T   K   S   P   V   F   G   M   I   N   S   T   I   W   T  960

WT  ctc tcc acc ata aga agt tct aac tct cag gac cga ctt ctt aac tca ttt gac gat gca 2940
WT   L   S   T   I   R   S   S   N   S   Q   D   R   L   L   N   S   F   D   D   A  980
Res ctc cac cat aag aag ttc taa ctc tca gga ccg act tct taa ctc att tga cga tgc aca 2938
Res  L   H   H   K   K   F   -   L   S   G   P   T   S   -   L   I   -   R   C   T  980
```
