## Supplementary material for "Resistance bioassays and allele characterisation inform analysis of *Spodoptera frugiperda* (Lepidoptera: Noctuidae) introduction pathways in Asia and Australia": Fig. S15

**Fig. S15:** Alignment of the amino acid sequences of the potential ABCC2 frameshift mutation in *Spodoptera frugiperda* from Australia.

```
CLUSTAL O(1.2.4) multiple sequence alignment


ABCC2_SFRICE002622      MDKSNKNTAANGDGGQRAGEPKERVRKKPNILSRIFVWWIFPVLITGNKRDVEEDDLIVP	60
R_allele                MDKSNKNTAANGDGGQRAGEPKERVRKKPNILSRIFVWWIFPVLITGNKRDVEEDDLIVP	60
                        ************************************************************

ABCC2_SFRICE002622      SKKFNSERQGEYFERYWFEEVAIAEREDRDPSLWKAMRRAYWLQYMPGAIFVLLISGLRT	120
R_allele                SKKFNSERQGEYFERYWFEEVAIAEREDRDPSLWKAMRRAYWLQYMPGAIFVLLISGLRT	120
                        ************************************************************

ABCC2_SFRICE002622      AQPLLFSQLLSYWSVDSEMSQQDAGLYALAMLGINFITMMCTHHNNLFVMRFSMKVKIAA	180
R_allele                AQPLLFSQLLSYWSVDSEMSQQDAGLYALAMLGINFITMMCTHHNNLFVMRFSMKVKIAA	180
                        ************************************************************

ABCC2_SFRICE002622      SSLLFRKLLRMSQVSVGDVAGGKLVNLLSNDVARFDYAFMFLHYLWVVPLQVGVVLYFVY	240
R_allele                SSLLFRKLLRMSQVSVGDVAGGKLVNLLSNDVARFDYAFMFLHYLWVVPLQVGVVLYFVY	240
                        ************************************************************

ABCC2_SFRICE002622      DAAGWAPYVGLFGVIILIMPLQAGLTKLTGVVRRMTAKRTDKRIKLMSEIINGIQVIKMY	300
R_allele                DAAGWAPYVGLFGVIILIMPLQAGLTKLTGVVRRMTAKRTDKRIKLMSEIINGIQVIKMY	300
                        ************************************************************

ABCC2_SFRICE002622      AWEKPFQLVVKAARAYEMSALRKSIFIRSMFLGFMLFTERSVMFLTVLTLALTGNMISAT	360
R_allele                AWEKPFQLVVKAARAYEMSALRKSIFIRSMFLGFMLFTERSVMFLTVLTLALTGNMISAT	360
                        ************************************************************

ABCC2_SFRICE002622      LIYPIQQYFGIITMNVTLILPMAFASFSEMLISLERIQGFLLLDERSDIQITPKVVNGAG	420
R_allele                LIYPIQQYFGIITMNVTLILPMAFASFSEMLISLERIQGFLLLDERSDIQITPKVVNGAG	420
                        ************************************************************

ABCC2_SFRICE002622      SKLFNNSKKEGGLETGIVLPTKYSPTEANIARPMQDEPNMADYPVQLNKVNATWADLNDN	480
R_allele                SKLFNNSKKEGGLETGIVLPTKYSPTEANIARPMQDEPNMADYPVQLNKVNATWADLNDN	480
                        ************************************************************

ABCC2_SFRICE002622      KEMTLKNISLRVRKNKLCAVIGPVGSGKTSLLQLLLRELPVTSGNLSISGTVSYASQEPW	540
R_allele                KEMTLKNISLRVRKNKLCAVIGPVGSGKTSLLQLLLRELPVTSGNLSISGTVSYASQEPW	540
                        ************************************************************

ABCC2_SFRICE002622      LFPATVRENILFGLEYNVAKYKEVCKVCSLLPDFKQFPYGDLSLVGERGVSLSGGQRARI	600
R_allele                LFPATVRENILFGLEYNVAKYKEVCKVCSLLPDFKQFPYGDLSLVGERGVSLSGGQRARI	600
                        ************************************************************

ABCC2_SFRICE002622      NLARAVYREADIYLLDDPLSAVDANVGRQLFDGCIKGYLSGKTCILVTHQIHYLKAADFI	660
R_allele                NLARAVYREADIYLLDDPLSAVDANVGRQLFDGCIKGYLSGKTCILVTHQIHYLKAADFI	660
                        ************************************************************

ABCC2_SFRICE002622      VVLNEGSVENMGSYDELMKTGTEFSMLLSDQASEGSDTDKKERPAMMRGISKMSVKSDDE	720
R_allele                VVLNEGSVENMGSYDELMKTGTEFSMLLSDQASEGSDTDKKERPAMMRGISKMSVKSDDE	720
                        ************************************************************

ABCC2_SFRICE002622      EGEEKVQVLEAEERQSGSLKWDVLGRYMKSVNSWCMVVMAFLVLVITQGAATTTDYWLSF	780
R_allele                EGEEKVQVLEAEERQSGSLKWDVLGRYMKSVNSWCMVVMAFLVLVITQGAATTTDYWLSF	780
                        ************************************************************

ABCC2_SFRICE002622      WTNQVDGYIQTLPEGESPNPELNTQVGLLTTGQYLIVHGSVVLAIIILTQVRILSFVVMT	840
R_allele                WTNQVDGYIQTLPEGESPNPELNTQVGLLTTGQYLIVHGSVVLAIIILTQVRILSFVVMT	840
                        ************************************************************

ABCC2_SFRICE002622      MRASENLHNTIYEKLIVAVMRFFDTNPSGRVLNRFSKDMGAMDELLPRSMLETVQMYLSL	900
R_allele                MRASENLHNTIYEKLIVAVMRFFDTNPSGRVLNRFSKDMGAMDELLPRSMLETVQMYLSL	900
                        ************************************************************

ABCC2_SFRICE002622      ASVLVLNAIALPWTLIPTTVLMFIFVFLLKWYINAAQAVKRLEGTTKSPVFGMINSTISG	960
R_allele                ASVLVLNAIALPWTLIPTTVLMFIFVFLLKWYINAAQAVKRLEGTTKSPVFGMINSTIWT	960
                        **********************************************************

ABCC2_SFRICE002622      LSTIRSSNSQDRLLNSFDDAQNLHTSAFYTFLGGSTAFGLYLDTLCLIYLGIIMSIFILG	1020
R_allele                LHHKKF------------------------------------------------------	966
                        *   :

ABCC2_SFRICE002622      DFGELIPVGSVGLAVSQSMVLTMMLQMAAKFTADFLGQMTAVERVLEYTKLPTEENMETG	1080
R_allele                ------------------------------------------------------------	966


ABCC2_SFRICE002622      PTTPPKDWPSAGEVTFSNVYLKYSPDDPPVLKDLNFAIKSGWKVGVVGRTGAGKSSLISA	1140
R_allele                ------------------------------------------------------------	966


ABCC2_SFRICE002622      LFRLSDITGSIKIDGLDTQGIAKKLLRSKISIIPQEPVLFSASLRYNLDPFDNYNDDDIW	1200
R_allele                ------------------------------------------------------------	966


ABCC2_SFRICE002622      RALEQVELKESIPALDYKVSEGGTNFSMGQRQLVCLARAILRSNKILIMDEATANVDPQT	1260
R_allele                ------------------------------------------------------------	966


ABCC2_SFRICE002622      DALIQKTIRKQFATCTVLTIAHRLNTIMDSDRVLVMDQGVAAEFDHPYILLSNPNSKFSS	1320
R_allele                ------------------------------------------------------------	966


ABCC2_SFRICE002622      MVKETGDNMSRILFEVAKTKYESDSKTA	1348
R_allele                ----------------------------	966
```
